# Normal mammary epithelia accumulate DNA methylation changes with age that are consistent with breast cancer methylomes

**DOI:** 10.1101/2020.06.26.170043

**Authors:** Parijat Senapati, Masaru Miyano, Rosalyn W. Sayaman, Mudaser Basam, Candi Trac, Amy Leung, Mark A. LaBarge, Dustin E. Schones

## Abstract

Luminal epithelial cells (LEps) are a key cell lineage implicated in age-related luminal breast cancers. Alterations to the epigenome are a hallmark of aging cells. However, the extent of age-associated DNA methylation alterations in LEps, and the corresponding functional consequences of these alterations, have remained unclear. We report here that aging leads to distinct methylation changes in LEps. Luminal lineage-specific genes gain promoter methylation, whereas myoepithelial-specific genes lose promoter methylation. Regulatory elements display methylation changes at lineage-specific TF binding sites consistent with the loss of lineage fidelity. CpG islands (CGIs) and transposable elements (TEs) have stochastic methylation gain and loss, respectively. PRC2 target genes that are hypermethylated in luminal breast cancer exhibit stochastic methylation increase with age. TEs with stochastic methylation loss are activated in breast cancer and potentially function as regulatory elements contributing to the loss of lineage fidelity with age. Each of these classes of methylation changes impact the regulation of genes associated with luminal breast cancer. Altogether, our results indicate that aging leads to DNA methylation changes that could determine breast cancer susceptibility.

**Significance:** Mammary luminal epithelial cells lose lineage-specific expression with age and accumulate stochastic methylation changes that lead to loss of silencing at transposable elements. These age-dependent events potentially promote breast cancer susceptibility.

## Introduction

Over 75% of breast cancer diagnoses are in women over the age of 50, making age the most significant risk factor for breast cancer (1). This is especially true for luminal subtypes of all age-related breast cancers (2). The mammary epithelium is composed of two principal lineages, myoepithelial cells (MEps) and luminal epithelial cells (LEps), with the latter being the chief suspect cell of origin for luminal breast cancers (3,4). Most breast cancers are sporadic with no known inherited mutation or family history. A better understanding of how age-dependent molecular changes contribute to breast cancer risk is critical for designing better therapies and prevention strategies.

Aging in the breast tissue is associated with significant physiological changes that lead to changes in molecular properties and the proportion of the different cell populations with age (5-8). Human mammary epithelial cells (HMECs) from older post-menopausal women, when transformed *in vitro*, exhibit gene expression patterns characteristic of luminal breast cancer subtypes. In contrast, transformed HMECs from younger pre-menopausal women show more properties of basal breast cancers (9). Striking age-dependent changes in abundance and phenotype occur in LEps, which gain expression of characteristic MEp genes (6,10,11) and signaling molecules consistent with those of immortalized luminal cells (10), suggesting that age-associated epigenetic changes in luminal cells are directly related to elevated cancer risk.

DNA methylation is a relatively stable component of the epigenome that gets established in early development and is maintained through successive cycles of cell division to preserve cell identity (12). DNA methylation patterns, however, are known to change with age (13–16) and DNA methylation levels of specific CpG sites show a significant correlation with chronological age (14). This ‘epigenetic clock’ can be used to predict the biological age of diverse human tissues (14). However, this pan-tissue epigenetic clock is “poorly calibrated” in breast tissue, even for women without cancer, possibly due to physiological factors such as changes in hormonal status with age (14). Previous studies examining DNA methylation changes in breast tissue with age were either performed in bulk mammary tissue with heterogeneous cell populations or interrogated only a subset of CpGs in the genome (17,18). A complete understanding of age-associated DNA methylation changes in human mammary LEps and how it relates to age-related breast cancer has been lacking.

We performed whole-genome bisulfite sequencing (WGBS) and Assay for Transposase-Accessible Chromatin (ATAC-seq) in non-senescent primary human mammary luminal epithelial cells from pre-menopausal (young) and post-menopausal (old) women. We find that LEps from older women display DNA methylation changes at regulatory elements associated with altered expression and activity of lineage-specific transcription factors such as ELF5 and TP63. We used chromatin state predictions from mammary epithelial cell lineages to show that MEp lineage-specific regulatory elements lose methylation, whereas luminal-specific regions gain methylation. Additionally, PRC2 (polycomb repressive complex 2) regulated genes gain DNA methylation at their promoters with age, consistent with these genes having methylated promoters in luminal breast cancer (19). We also observed increased methylation disorder with age, resulting in stochastic methylation gain at CpG islands (CGIs) and loss at evolutionarily recent transposable elements (TEs). Using breast cancer WGBS data, we show that TEs that lose methylation with age were also hypomethylated in breast cancers. In addition, we used ATAC-seq and RNA-seq data from the TCGA-BRCA cohort and RNA-seq done on DCIS (ductal carcinoma in situ) samples to show that TEs with age-dependent methylation loss are activated in breast cancer and can function as regulatory elements for nearby genes. Overall, our results suggest that aging mammary luminal epithelial cells undergo DNA methylation alterations at regulatory elements, CGIs, and evolutionarily recent TEs, similar to methylation changes observed in luminal breast cancer.

## Results

### Aging leads to altered DNA methylation patterns in mammary luminal epithelial cells

We performed whole-genome bisulfite sequencing (WGBS-seq) on human mammary LEps from younger (<30y) and older (>55y) women (**Figure S1A, Table S1**). The age groups were chosen to represent samples from women who were clearly pre-menopausal or post-menopausal to avoid confounding issues of peri-menopause. We previously demonstrated that LEps from these primary cultures maintain gene expression and DNA methylation profiles that mirror those in tissue (8,11). Moreover, these are non-senescent cells and therefore are a valuable resource that can be used to interrogate aging-dependent changes in the breast tissue – distinct from those arising from cellular senescence.

WGBS libraries were sequenced to ~30X genome coverage, generating ~ 1B 150-bp paired-end reads (~86 Gb uniquely mapped sequence per sample) (**Table S2**) to obtain a representation of about 95% of individual CpG sites in our samples (see Methods and **Tables S2, S3**). We compared our DNA methylation profiles with previously published WGBS studies on distinct human mammary epithelial cell types from reduction mammoplasty (RM) tissue of a young (19y old) donor (20) and found that the cultured LEps resemble the ER-negative (ER-) LEp population (**Figure S1B**). Young and old LEPs clustered separately indicating differences in methylomes (**Figure S1B**) that were not apparent when considering total methylation levels (**Figure S1C, D**). Pairwise comparisons of methylomes also showed higher number of differentially methylated regions (DMRs) between young and old pairs (**Figure S1E**). Comparing the methylomes of old versus young LEps, we identified 18,860 differentially methylated regions (DMRs) (see Methods), with 7,207 showing methylation loss and 11,653 showing methylation gain in old LEps. Most DMRs were 100-500bp in length (**Figure S1F**). The methylation changes at DMRs were confirmed with heatmaps showing methylation levels across samples (**Figure 1A**), as well as kernel smoothed methylation plots of representative DMRs (**Figure 1B**). The majority of DNA methylation changes were observed at non-promoter regions (**Figure S1G-H**).

**Figure 1:**
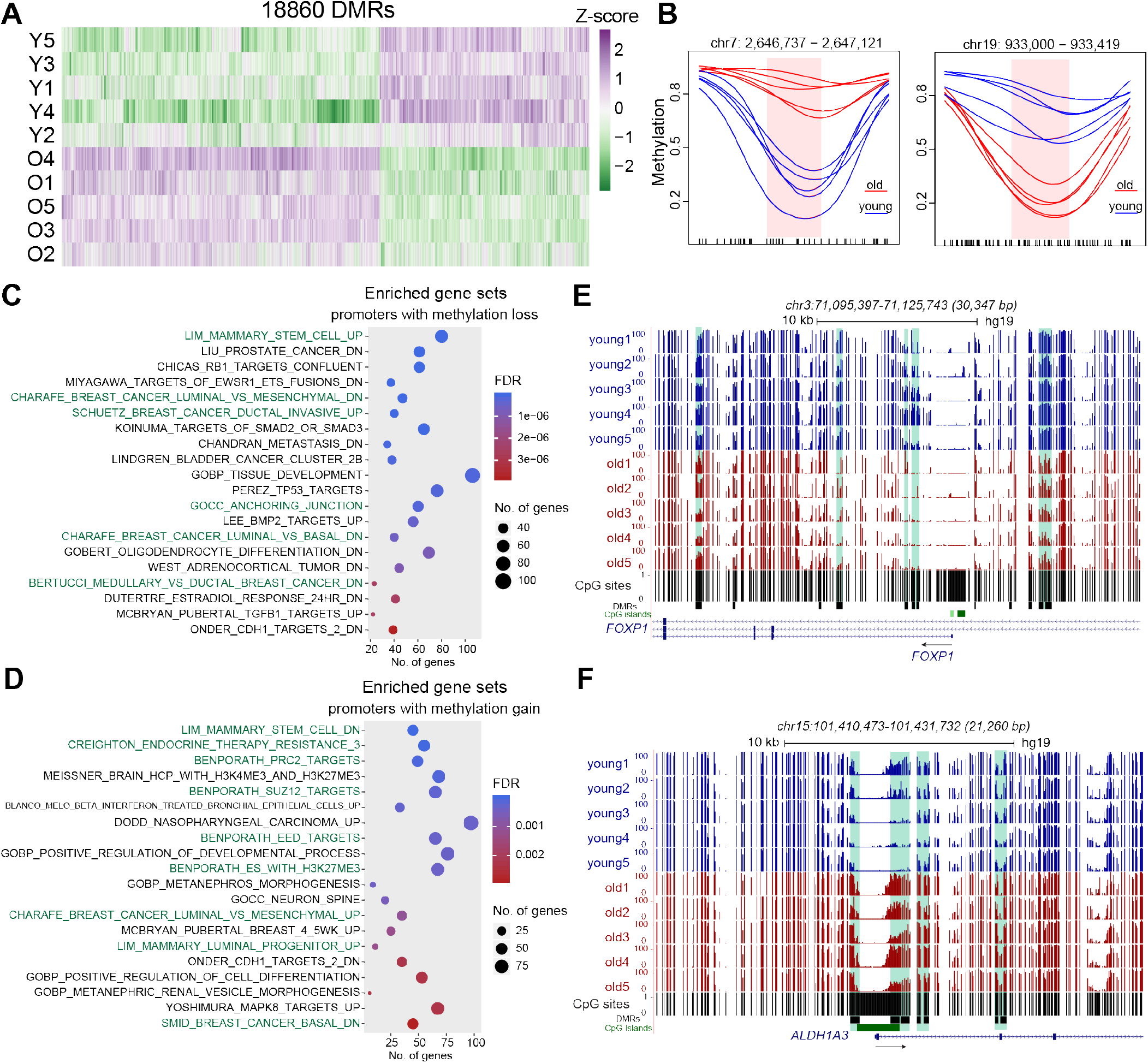
Mammary luminal epithelial cells from older individuals show DNA methylation changes at distinct genomic loci. (A) Methylation levels of identified DMRs across all samples. Samples are in rows, and DMRs are in columns. The color intensity represents the Z-score scaled from the mean. Values ranging from higher to lower methylation are shown as purple to green. (B) Kernel smoothed line plots of representative DMRs±500bp with methylation gain (left panel) and methylation loss (right panel). Blue and red lines respectively represent young and old samples. DMRs are highlighted in pink and black ticks mark each CpG position. (C) MSigDB gene sets enriched in genes with age dependent DNA methylation loss at promoters (TSS+/-2kb). (D) MSigDB gene sets enriched in genes with age dependent DNA methylation gain at promoters (TSS+/-2kb). Relevant gene sets are highlighted in green. C-D. UP: upregulated, DN: downregulated. (E) Genome browser tracks showing DMRs with methylation loss in old LEps at the *FOXP1* gene locus. (F) Genome browser tracks showing methylation gain at the CpG island promoter of *ALDH1A3* gene. E-F. DMR positions are shown as black boxes.

We assigned DMRs to genes based on overlap with either promoters or gene body (exonic or intronic) (**Table S4A-B**). We used the Molecular Signatures Database (MSigDB) to identify the gene sets or pathways over-represented among these genes (**Table S5A-D**). Genes with methylation loss at promoters belong to gene sets expressed in mammary stem cells, cell-cell anchoring junctions epithelial-to-mesenchymal transition, and genes downregulated in luminal breast cancer (**Figure 1C, Table S5A**). Genes with promoter methylation gain were enriched for luminal specific genes upregulated in luminal breast cancer and Polycomb (PRC2) target genes (**Figure 1D, Table S5C**, BENPORATH gene sets). We also noted that genes expressed in mammary stem cell enriched basal compartment showed methylation loss (**Figure S2A**) whereas luminal lineage restricted genes showed methylation gain (**Figure S2B**). As an example, *FOXP1*, a mammary stem cell specific gene essential for differentiation into various mammary epithelial lineages (21), shows methylation loss at several sites its promoter and gene body (**Figure 1E**). *ALDH1A3*, on the other hand, is an ER-luminal lineage restricted gene (22) which gains methylation at several sites including the promoter in old LEps (**Figure 1F**). DNA methylation changes with age suggest that luminal cells increase stem cell related properties while repressing genes that are specifically related to maintenance of the luminal lineage.

### Aging-associated DNA methylation changes occur at lineage-specific transcription factor binding sites

To investigate whether DNA methylation changes were associated with transcription factor (TF) binding signatures, we performed motif enrichment analysis (see Supplemental Methods for details) using position weight matrices (PWMs) from the JASPAR database (23). DMRs that lose methylation are enriched for binding sites of TFs such as EGR1, TP63, TP73, TEAD1, TEAD4, and RUNX1 (**Figure 2A left panel, Table S6A**). Several of these TF binding sites, including TEAD, TP63, TP73, NFI, and EGR1, are enriched at active regulatory elements in basal/myoepithelial cells (20). To investigate whether these TFs change expression with age, we checked their expression using RNA-sequencing data from LEps and MEps of young and old individuals (11). *TP63* and *NFIA (NFI family TF; binds to NFI half-sites)* showed increased expression in old compared to young LEps (**Figure 2B and S3A**). We also checked whether these TFs showed any DNA methylation changes. *EGR1* showed significant methylation loss at its promoter (**Figure S3B**) whereas *NFIA* and *TP63* exhibit methylation loss at specific sites within their gene bodies (**Table S4A**) suggesting altered methylation at regulatory elements. Although not all TFs showed an increase in expression with age, we noted that *TP63, NFIA, EGR1, and TP73* have relatively higher expression in MEps as compared to LEps (**Figure 2A right panel and 2E**). The expression patterns of TFs and the hypomethylation of their targets are concordant with the known roles of these TFs. For example, young LEps do not express *TP63*, which is a MEp lineage-specific transcription factor (8). Furthermore, TEAD1 and TEAD4 are DNA binding partners for YAP (Yes-associated protein), a transcriptional co-activator regulated by the Hippo signaling pathway. YAP, typically a MEp specific protein, is sequestered in the cytoplasm through activity of kinases downstream of the Hippo pathway (24). Old LEps exhibit increased *TP63* expression and nuclear YAP localization (7,8), which leads to increased TEAD binding and YAP activity. YAP is also known to bind p73 (TP73 protein) and RUNX2 (24). These results suggest that increased expression/activity of myoepithelial-specific TFs in old LEps lead to increased binding and loss of DNA methylation in these regions (**Figure 2F**).

**Figure 2:**
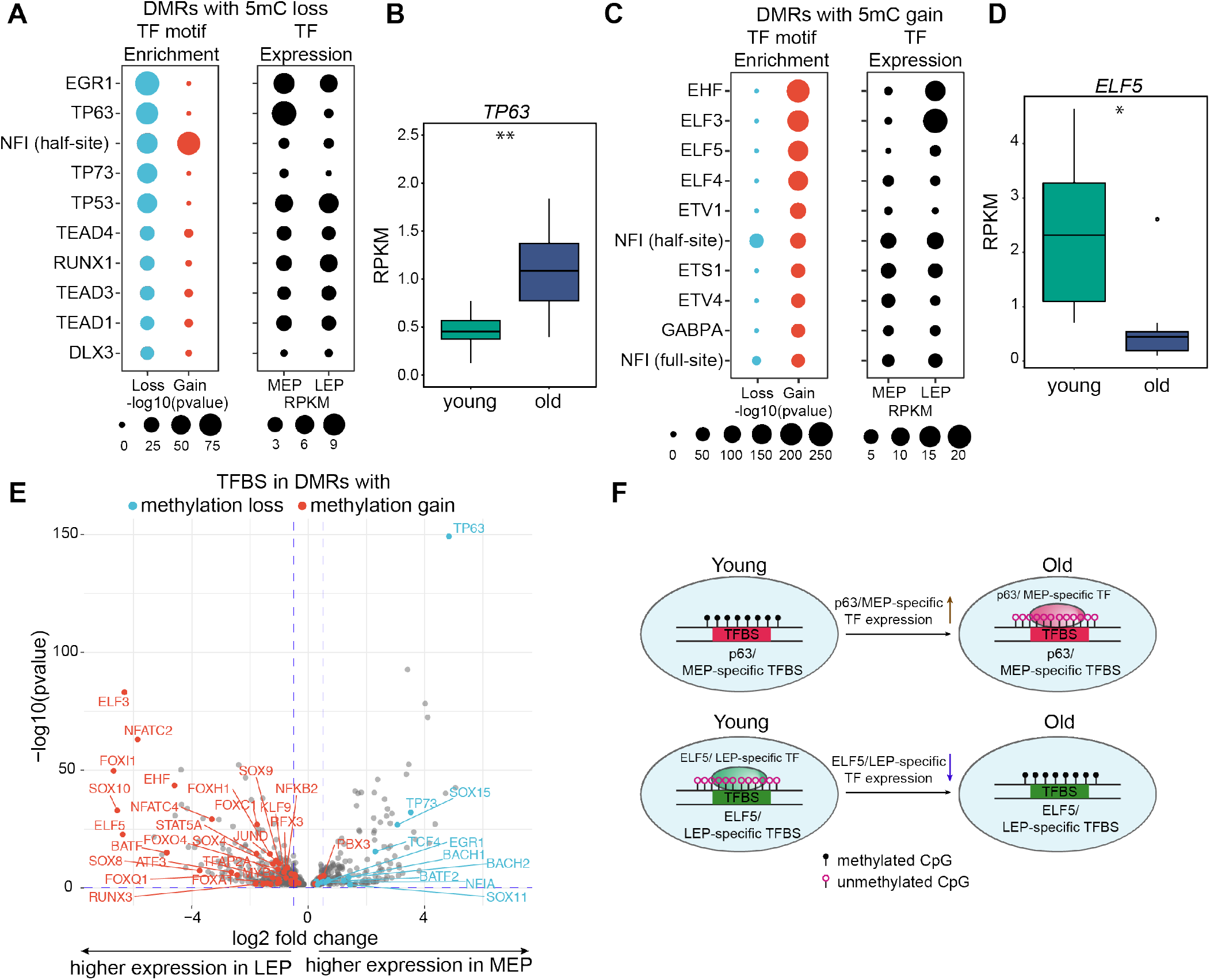
Differentially methylated regions show enrichment of lineage-specific transcription factor binding sites. (A) Top ten TF binding sites enriched in the DMRs that lose methylation with age are shown. Dots in the left panel show -log10(p-value) for the enrichment of the transcription factor binding site. Dots in the right panel show expression levels (RPKM) of the respective transcription factor in young myoepithelial (MEp) and luminal (LEp) cells. (B) Boxplot showing expression levels of myoepithelial-specific TF TP63 in young and old LEps (n=8, *p*-value < 0.01; two-tailed unpaired t test with Welch’s correction). (C) Top ten TF binding sites enriched in the DMRs that gain methylation with age are shown. Dots in the left panel show -log10(p-value) for the enrichment of the TF binding site. Dots in the right panel show expression levels (RPKM) of the respective transcription factor in young myoepithelial (MEp) and luminal (LEp) cells. (D) Boxplot showing expression levels of luminal lineage-specific TF ELF5 in young and old LEps (n=8, *p*-value < 0.05; two-tailed Mann Whitney test). (E) Volcano plot showing expression levels of all TFs expressed in LEPs and MEPs (RPKM > 0.1). TFs with motifs enriched in DMRs with methylation loss and gain are highlighted in blue and red respectively. *P*-value < 0.05. (F) Schematic showing the role of myoepithelial-specific transcription factors in causing loss of methylation. Conversely, loss of expression of luminal-specific transcription factors leads to methylation of unbound sites and methylation gain.

DMRs with methylation gain are enriched for binding sites of the ETS family of transcription factors EHF, ELF3, ELF5, ELF4, ETV1, ETS1, ETV4, GABPA, and NFI half and full sites (**Figure 2C left panel, Table S6B**). ELF5 is a luminal lineage-specific transcription factor that decreases expression with age (**Figure 2D**) (8). Both *ELF3* and *ELF5* genes also show significant methylation gain at their promoter and gene locus (**Figure S3C-D**). In addition, *ELF5, EHF* and *ELF3* exhibit relatively higher expression in LEps as compared to MEps (**Figure 2C right panel and 2E**). These results further suggest that decreased expression of luminal lineage-specific TFs may lead to a decrease in binding and protection of binding sites from DNA methyltransferases (DNMTs). This, in turn, may result in the methylation and repression of these luminal lineage-specific regions (**Figure 2F**). We also examined the known DNA methylation binding sensitivities of TFs (25) with enriched binding sites. TFs with binding sites showing methylation loss either do not have a CpG at their respective site or their binding is unaffected by DNA methylation (**Table S7**). Presumably they recruit other factors such as TETs to bring about DNA methylation changes. However, TFs with binding sites showing methylation gain are sensitive to DNA methylation (**Table S7**). Generally, sites of methylation loss were enriched for myoepithelial-specific TF binding motifs (**Figure 2E**), whereas sites of methylation gain were enriched for luminal lineage-specific TF binding motifs (**Figure 2E**). For example, SOX4 sites are enriched in DMRs with methylation gain, and it is a luminal-specific TF (**Figure 2E**). SOX15, on the other hand, is a myoepithelial-specific TF and its binding sites are enriched at DMRs with methylation loss (**Figure 2E**). Expression of *SOX4* decreases, whereas *SOX15* increases with age in old LEps (**Figure S3E-F**), indicating loss of expression of luminal lineage-specific TFs and increased expression of myoepithelial-specific TFs. Collectively, we observe that aging-associated DNA methylation changes in LEps occur at lineage-specific TF binding sites that seem to contribute to loss of cell identity. Enticingly this is a characteristic of cancer cells whereby even the so-called luminal subtype cancer cells exhibit expression of genes and proteins that are characteristic of basal cells.

### Lineage-specific regulatory regions are hotspots for methylation changes with age

Given that aging-associated DNA methylation changes occurred at regions containing lineage-specific TF binding sites, we examined whether these binding sites are functional by measuring chromatin accessibility using ATAC-seq in young and old LEps (**Table S1**). Open chromatin region at a TF binding site is a reliable indicator of a regulatory element with bound TFs as not all TF binding sites in a given condition in a cell are bound. DMRs with methylation loss showed corresponding increased accessibility in older LEps (**Figure 3A-B**). On the other hand, DMRs with methylation gain showed a decrease in chromatin accessibility (**Figure 3C-D**). These results indicate that functional regulatory elements undergo DNA methylation changes with age.

**Figure 3:**
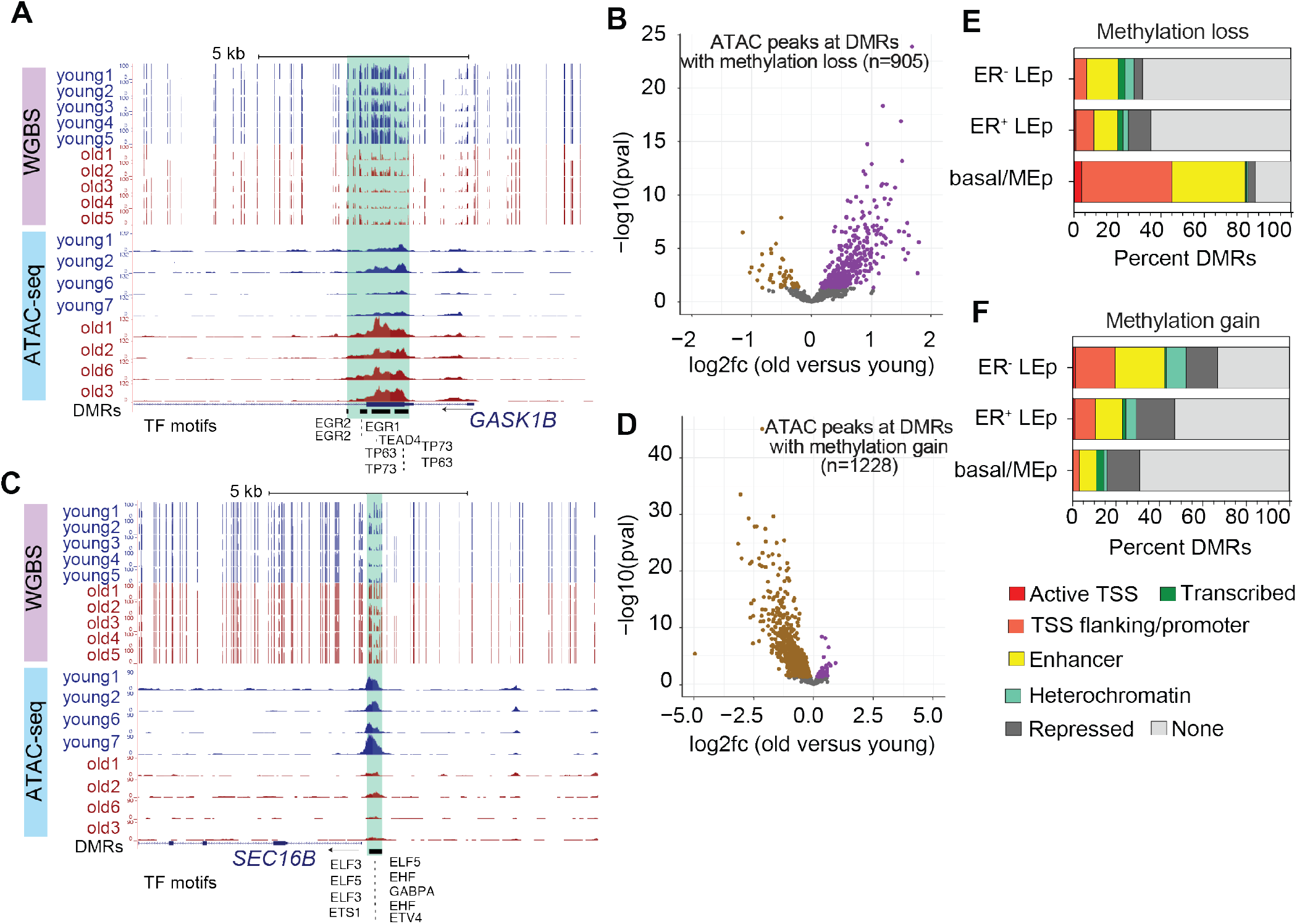
Differentially methylated regions are enriched at lineage-specific regulatory elements. (A) Genome browser tracks showing a representative example of a DMR with methylation loss accompanied with increase in chromatin accessibility in old LEPs. DMRs are highlighted in green. Bottom tracks show location of MEp-specific TF binding sites. (B) Volcano plot showing log2 fold change (log2fc) of ATAC-seq peaks overlapping DMRs with methylation loss in old versus young LEPs. Positive log2 fold change indicates increase in accessibility. (C) Genome browser tracks showing a representative example of a DMR with methylation gain accompanied with decrease in chromatin accessibility in old LEPs. DMRs are highlighted in green. Bottom tracks show location of LEp-specific TF binding sites. (D) Volcano plot showing log2 fold change (log2fc) of ATAC-seq peaks overlapping DMRs with methylation gain in old versus young LEPs. Negative log2 fold change indicates decrease in accessibility. (E) Percentage number of DMRs with methylation loss and (F) methylation gain that overlap with chromHMM features of ER^-^, ER^+^ luminal cells, and basal cells (rows) is shown. The color key indicates the various chromatin states identified from ChIP-seq datasets.

To further investigate the functional relevance of the age-dependent DNA methylation changes in the breast tissue, we used published histone modification ChIP-seq datasets in mammary epithelial lineages to predict chromatin states at DMRs (20). To discern whether age-dependent DNA methylation changes were occurring at lineage-specific regulatory elements, we predicted chromatin states in ER^+^ luminal, ER^-^ luminal, and basal (MEp) cells from RM tissue of a young (19y old) donor (20). We classified DMRs into seven different chromatin states that are characteristic of i) Active TSS, ii) TSS flanking/promoter, iii) transcribed, iv) Enhancer, v) Heterochromatin, vi) Repressed and vii) None/unclassified regions based on the combinatorial histone modification levels in each cell type (see Supplemental Methods; **Figure S4A-C**). DMRs with methylation loss in old LEps overlap primarily with MEp-specific promoter and enhancer regions (~80% of methylation loss DMRs) (**Figure 3E**). Because these sites also contain binding sites for TFs enriched at MEp-specific active enhancers (20), these results indicate that MEp-specific regulatory elements lose methylation in old LEps. Active enhancer sites are hypomethylated (26,27) and methylation loss at MEp-specific regulatory elements might indicate an epigenetic shift towards MEp gene expression patterns in LEps with age.

Conversely, DMRs with methylation gain overlap putative promoter and enhancer regions in ER^-^ luminal cells, accounting for about 42% of methylation gain DMRs (**Figure 3F**). As these DMRs also contain binding sites for TFs enriched at luminal lineage-specific active enhancers (20), these results indicate that luminal-specific regulatory regions gain DNA methylation in aging LEps. Together, these results suggest that aging LEps acquire permissive chromatin states at MEp-specific regions and repress ER^-^ luminal-specific chromatin regions through changes in DNA methylation.

### Polycomb repressive complex 2 target genes gain DNA methylation with age

The Polycomb repressive complex 2 (PRC2) is a histone methyltransferase complex that trimethylates lysine 27 of histone H3 (H3K27me3) and mediates transcriptional silencing of developmentally important genes. As per current understanding PRC2 is targeted to unmethylated CpG-dense regions, primarily CpG islands (CGIs) that occur at gene promoters (28). Earlier we found that genes with methylation gain at their promoters were enriched in PRC2 target genes identified in embryonic stem cells (ESCs) (**Figure 1D**). Moreover, about 15% of the DMRs with methylation gain overlapped with Polycomb repressed regions marked by H3K27me3 (**Figure 3F and S4A-C**). PRC2 target gene promoters have been described to gain DNA methylation with age in various tissues as well as in mice (13,29-32). They are also known to be hypermethylated in diverse types of cancer, some among them are known tumor suppressor genes (33). Since the MSigDB PRC2 target gene sets are ESC-specific, we used H3K27me3 ChIP-seq data from ER^-^ and ER^+^ luminal cells, and basal cells (20) to determine if genes with promoter methylation gain are PRC2 targets in mammary epithelial lineages. We found that 39-46% of promoters that gain methylation with age are marked by H3K27me3 across the three cell lineages (**Figure S5A**). In contrast, only 14-27% of promoters with methylation loss are enriched for H3K27me3 (**Figure S5B**). Promoters with DNA methylation gain also have conserved H3K27me3 occupancy in all three epithelial lineages (**Figure S5C**) as compared to promoters with DNA methylation loss that show H3K27me3 only in the luminal lineage (**Figure S5D**). These results indicate that promoters with age-dependent DNA methylation gain are indeed *bona fide* PRC2 target genes in mammary epithelial cells. These results are consistent with PRC2 protecting unmethylated CpG-rich regions and that erosion of the PRC2 targeting with aging (32) may be fundamentally related to increased methylation at its target genes.

### Stochastic methylation loss at transposable elements (TE) and methylation gain at CpG islands (CGIs) with age

Aging is associated with increased transcriptional variability, cell-to-cell heterogeneity, and DNA methylation variability (34–37). Because we use a sorted luminal lineage cell population, we can discern DNA methylation variability in LEps eliminating confounding problems such as changes in cell proportions with age. To examine whether there is an increase in methylation heterogeneity in older individuals, we calculated methylation entropy, a measure of methylation heterogeneity between adjacent CpGs, for 150 bp non-overlapping windows across the genome (38). There are two modes by which a given region could increase methylation entropy. An unmethylated region can gain methylation stochastically or a methylated region can lose methylation stochastically (**Figure 4A-B**). Considering these regions, we observed an overall increase in methylation entropy in the old LEps. We identified regions with greater methylation entropy in older individuals and found a net decrease in methylation levels at these regions (**Figure S6A-D**), consistent with the fact that most of the genome is methylated in normal cells. Of these, unmethylated regions with increased methylation entropy were mainly at CGIs (69.2%; 1017 of 1469; **Figure 4C**). Conversely, methylated regions with increased entropy were found mainly at repeat regions (70.3%; 76575 of 108877; **Figure 4D**), which are known to be enriched with TEs. These results indicate that increased DNA methylation heterogeneity with age leads to stochastic methylation gain at CGIs and methylation loss at repeat regions.

**Figure 4:**
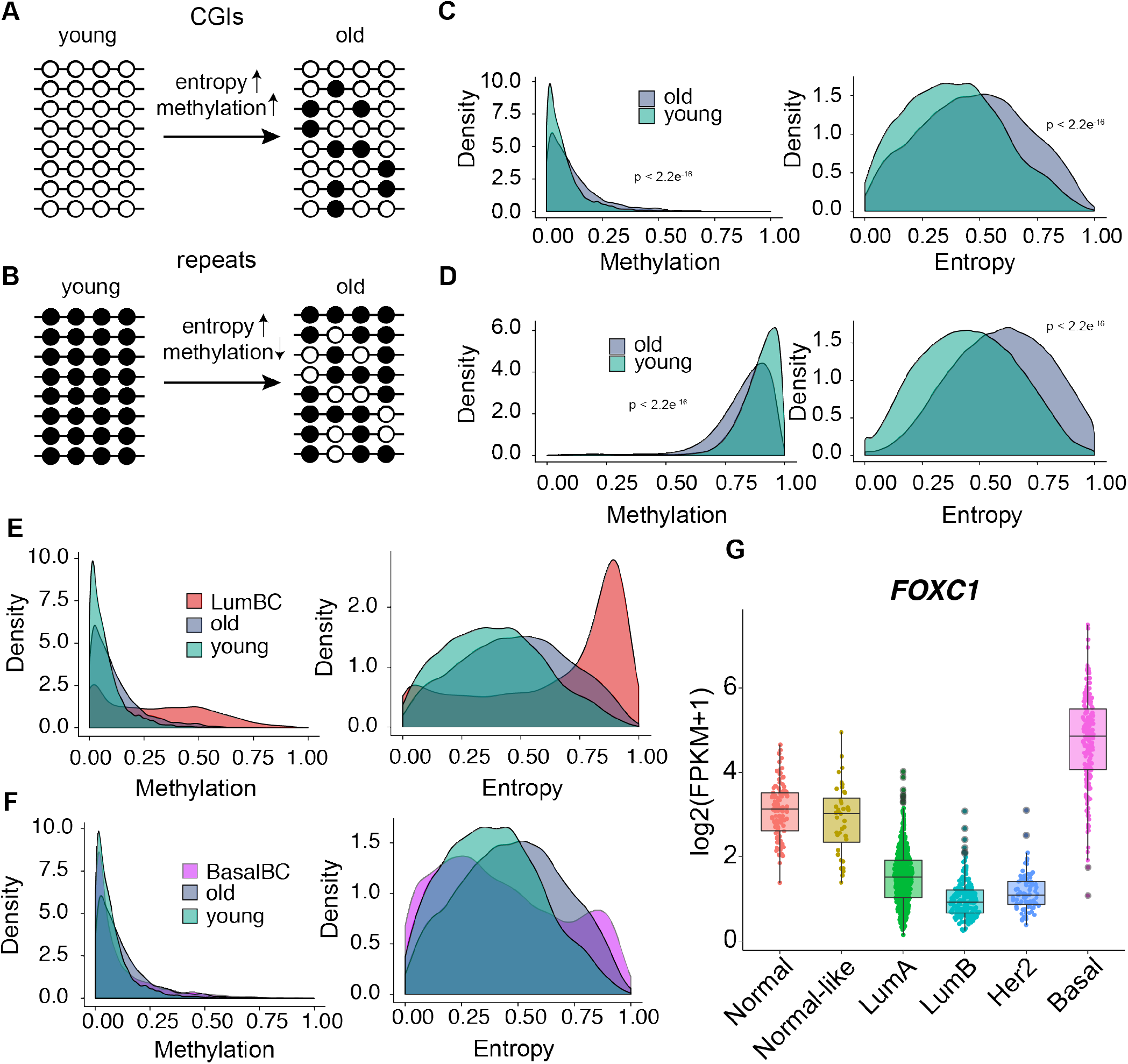
Aging leads to stochastic methylation loss at transposable elements and methylation gain at CpG islands (CGIs). (A) Schematic depicting changes in entropy associated with stochastic methylation gain at CGIs. (B) Schematic depicting changes in entropy associated with stochastic methylation loss at fully methylated regions such as repeat elements. (C) Methylation and entropy levels at unmethylated CGIs that show increase in entropy. (D) Alu methylation and entropy levels at fully methylated Alu elements that show increase in entropy. C-D. Significant differences in distributions were tested using Kolmogorov–Smirnov test. (E) Aggregate profiles of methylation and entropy levels at unmethylated CGIs that show increase in entropy with age. Luminal breast cancer samples (red) show an increase in DNA methylation as well as entropy at these CGIs. (F) Aggregate profiles of methylation and entropy levels at unmethylated CGIs that show increase in entropy with age. Basal breast cancer samples (pink) do not show an increase in DNA methylation or entropy at these CGIs. (G) Expression of *FOXC1* in normal tissues and five PAM50 subtypes of breast tumors from the TCGA BRCA collection. Luminal A (LumA) versus Basal p-value < 2.2×10^-16^; using two-tailed Wilcoxon signed-rank test. Luminal B (LumB) versus Basal (p-value < 2.2×10^-16^; using two-tailed Wilcoxon signed-rank test).

To examine the potential of stochastic methylation changes to have consequences for breast cancer, we used a published WGBS dataset of 30 primary tumors containing ER+ (luminal breast cancer) and ER-(basal breast cancer) samples (39). At CGIs with increased entropy, luminal breast tumors showed an even greater increase in methylation and entropy than observed in old LEps (**Figure 4E**). In contrast, basal tumors did not show an increase in CGI methylation or entropy (**Figure 4F**), indicating that this behavior is specific to the luminal subtypes of breast cancer. We identified the subset of entropy-gained CGIs that were hypermethylated in at least 20% (n >= 6) of the breast tumors. These CGIs were enriched at PRC2 target genes, which are known to be hypermethylated in luminal breast cancer (19) and other types of cancer (**Figure S6E**). Of note, PRC2 target genes are repressed in normal tissues through H3K27me3, and although they show DNA methylation gain at their CGI promoters, expression changes are not expected. Apart from PRC2 target genes, we also found gene sets that are upregulated in basal breast cancer and downregulated in luminal B breast cancer to be enriched at these CGIs (**Figure S6E**). For example, *FOXC1* gene is expressed in LEps and normal breast tissue but downregulated specifically in luminal and Her2-enriched breast cancers (**Figure 4G**). Taken together, these results indicate that increased DNA methylation heterogeneity with age leads to stochastic methylation gain at CGIs which is further exacerbated in luminal breast cancer.

### Increased DNA methylation variability between old individuals at TEs

We next tested whether aging would lead to increased interindividual variability in addition to increased methylation entropy. We indeed do observe increased variability of CpG methylation in older individuals (**Figure S7A**). Since some amount of DNA methylation variability is expected in human populations and differences at individual CpG sites may not have functional consequences, we first identified systemic regions of variable methylation across all 10 individuals using the Systemic Interindividual Variation (SIV) approach (40) (see Supplemental Methods). We identified 23,921 regions that were variably methylated only in old samples and 25,585 regions that were specific to the young samples (**Figure S7B**). Regions variably methylated in old samples mainly showed methylation loss compared methylation levels in younger individuals (17,780 of 23,921; 74.3%). The variable regions were ~150p-450bp in length with some spanning 2-2.5kb (**Figure S7C and S7D**). We next examined whether variable regions contained TEs and found that 68.7% of the regions variable in young samples contained TEs as compared to 67.59% of control non-variable regions. 71.4% regions variable in old samples overlapped with TEs as compared to 67.65% of control non-variable regions (**Figure S7E**). Among the TEs that are variably methylated in old samples, about 76.9% lose methylation and 23.13% gain methylation (**Figure S7F**). We also confirmed that variable methylation at TEs was not due to mapping issues (**Figure S7G**).

One of the main functions of DNA methylation is to silence TEs, especially those that are evolutionarily recent and more likely to be capable of activity (41,42). To investigate whether the TEs that are variably methylated in old samples are evolutionarily recent, we calculated the Jukes-Cantor evolutionary age of each TE copy. Among the variably methylated TEs, the evolutionarily recent elements (inserted <100 million years ago; (43)) tended towards DNA methylation loss, whereas the older TE elements (>100 million years) tended to gain methylation in old cells (**Figure 5A-D**). These results indicate that evolutionarily recent TEs lose methylation with age in a stochastic manner. While evolutionarily recent Alu elements such as AluY and AluS are highly abundant in the genome, a few AluS subfamilies such as AluSz and AluSx were enriched above background for stochastic loss of methylation (**Figure 5E, S7H**). Other evolutionarily recent TE subfamilies were enriched at regions with variable methylation loss as well, including both L1PA (primate-specific LINE-1) and HERVH elements (**Figure 5E-F**).

**Figure 5:**
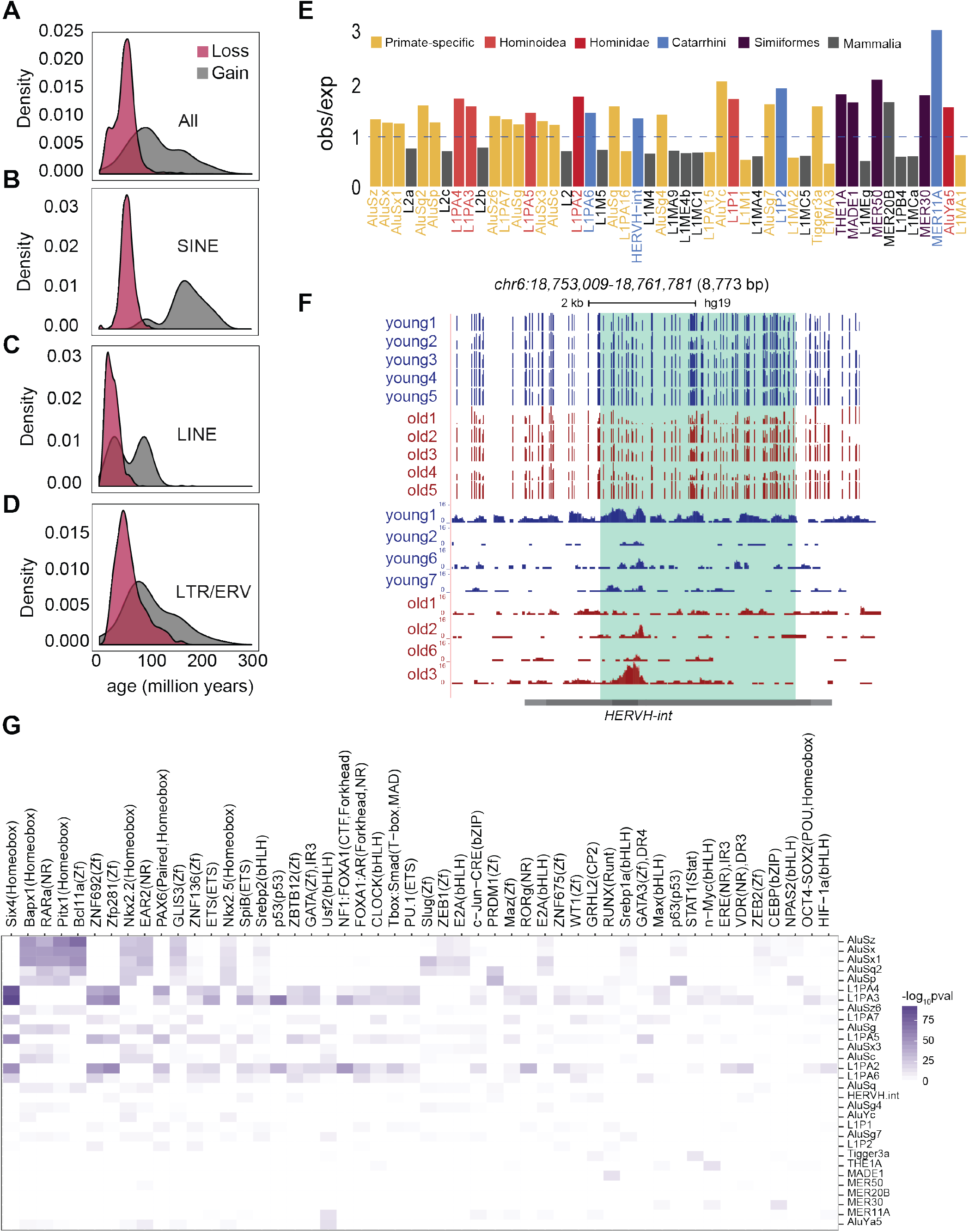
Variably methylated regions with methylation loss are enriched at primate-specific transposable elements. (A-D) Evolutionary age of (A) all TE elements (B) SINE, (C) LINE, and (D) LTR/ERV elements that are variably methylated with methylation loss or gain. Evolutionary ages of only the significantly enriched TE subfamilies are plotted. (E) TE subfamilies enriched in regions with variable methylation loss. Bars show observed/expected ratio for each TE subfamily. Expected number of overlaps in shuffled controls was used to calculate the ratio. Colors in the observed/expected graph highlight primate-specific or younger elements and grey bars indicate older elements. (F) Genome browser tracks showing a representative example of a HERVH-int element showing variable methylation loss in old samples. Bottom tracks are from ATAC-seq data showing gain of accessibility at the variably methylated region in the corresponding sample. (G) TF binding motifs enriched within each TE subfamily with variable methylation loss. Color scale is –log10(p-value) of TF motif enrichment.

Many TEs in the human genome have retained latent regulatory potential and can serve as regulatory elements when not suppressed (44–46). We therefore examined the TF motifs enriched at specific TE subfamilies with variable DNA methylation loss in old. We found that Alu subfamilies contain motifs for Homeobox TFs such as SIX4, BAPX1 and nuclear receptor TFs such as RARA which are upregulated in breast cancer (**Figure 5G**). Moreover, the younger L1PA subfamilies L1PA(2–4) have motifs for many TFs; the most relevant among them being binding sites for GATA, FOXA1 TFs and EREs which are bound by Estrogen receptor (ESR1) (**Figure 5G**). GATA3, FOXA1 and ESR1 are highly upregulated in luminal breast cancers (47). These results indicate that stochastic loss of methylation at these TEs can lead to binding of TFs expressed during breast cancer development.

To examine whether TEs similarly contribute TF binding sites to DMR regions associated with loss of lineage fidelity, we identified the TEs that overlap DMRs. Although many DMRs overlap TEs, given the high number of copies in the genome only a few TE subfamilies, especially those of the LTR class, were enriched over background. Several LTR subfamilies, including HERVH, MER52A, MER52D and LTR27E were enriched at DMRs with methylation loss in old LEps (**Figure S8A**) while LTR10C, HERVE, and MER11D (**Figure S8B**) were enriched in DMRs with methylation gain. Among the TEs that lose methylation, HERVH elements showed the highest enrichment for TEAD and RUNX TF binding sites (**Figure S8C**), motifs that were also found in non-repetitive DMRs with methylation loss (**Figure 2A**) and known binding partners of the MEp specific YAP. TP63 binding sites, which were also found at DMRs with methylation loss (**Figure 2A**) were enriched at AluSx, AluSp, L2, and L3 elements. ETS family TF binding sites that gain methylation were found within MIRb, MIR subfamilies of SINE elements and L2, L2a-c LINE subfamilies (**Figure S8D**), similar to non-repetitive DMRs with methylation gain (**Figure 2C**). SOX family TF binding sites that also occur in DMRs with methylation gain (**Figure 2E**) were present in AluY, LTR10 and MER11D elements with methylation gain (**Figure S8D**). Most of these DMR-associated TEs are evolutionarily older, indicating that they have been co-opted by lineage-specific TFs and contribute to aging-dependent DNA methylation changes in LEps.

### Transposable elements that lose methylation with age are activated in breast cancer

Given our observations that evolutionarily recent transposons lose methylation stochastically with age and contain motifs for TFs overexpressed in breast cancer, we examined whether breast cancers show loss of epigenetic suppression at these elements. Loss of TE suppression can lead to aberrant transcription of TE-derived transcripts and alternative isoforms due to splicing with neighboring genes in cancer cells (48). To investigate the potential of this happening in breast cancer, we examined published RNA-seq data from high-grade ductal carcinoma in situ (DCIS), a form of pre-cancer (49). We used the SQuIRE pipeline (50) to measure gene and TE expression changes in DCIS as compared to normal tissues; 554 age-hypomethylated TEs showed higher expression in DCIS. Of these, 468 were variably hypomethylated with age and 86 were within DMRs with methylation loss. As with the loci with variable methylation loss in older samples, HERVH elements and L1PA2 elements were most common among the TEs that showed aberrant expression. A representative example of a HERVH element that loses methylation in aging LEps and breast tumors is shown (**Figure 6A**). Aberrant transcription from this HERVH element was observed only in DCIS samples (**Figure 6A**). We also found examples of exonization of TEs, where a MIRb element was included within the exon of a non-coding RNA DRAIC expressed in DCIS (**Figure S9A**). While aberrant expression of hypomethylated TEs in DCIS was clear, we did not observe breast cancer subtype specificity in the expression of these TEs.

**Figure 6:**
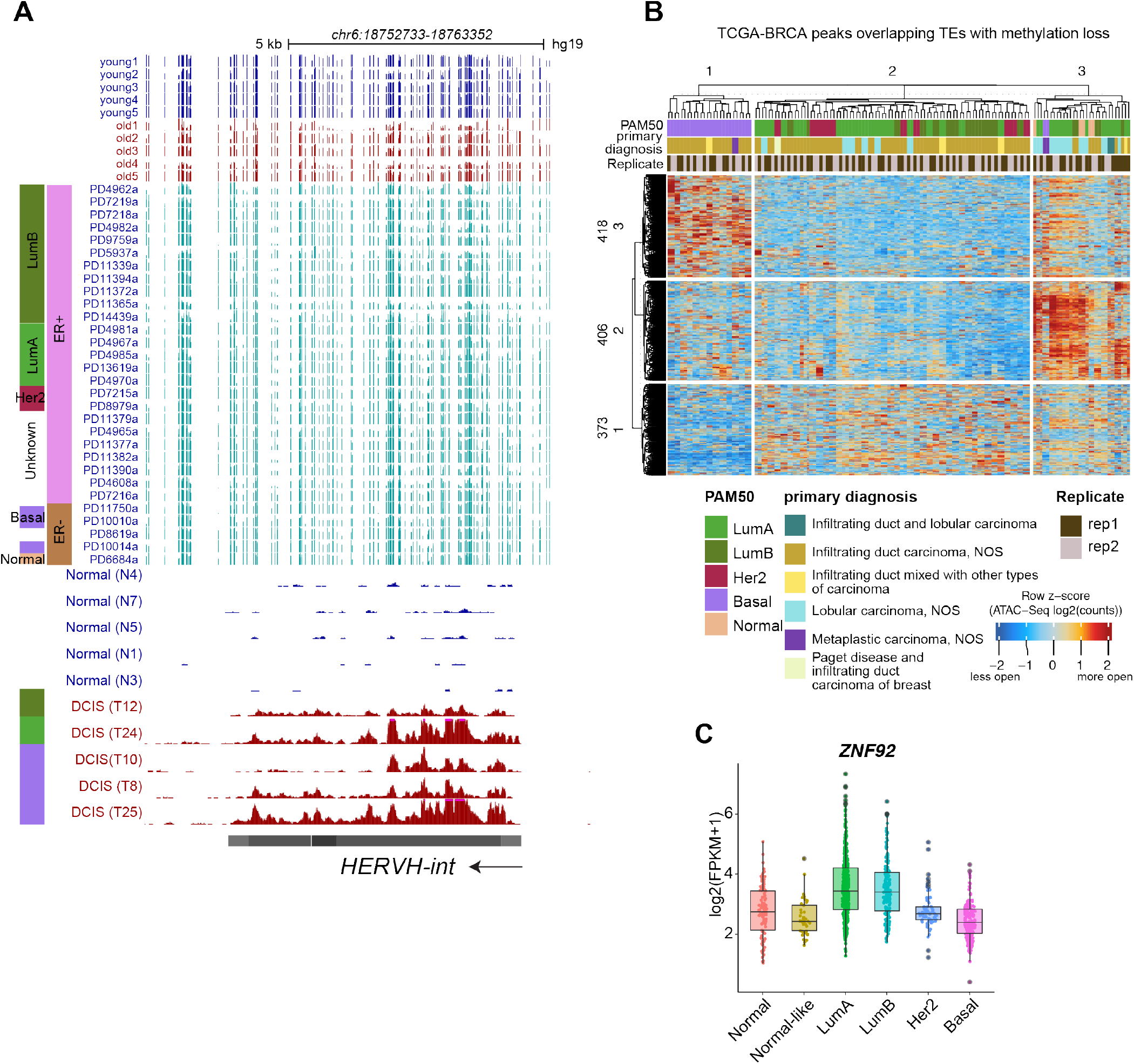
Transposable elements with age-dependent methylation loss are activated in breast cancer. (A) A representative HERVH element showing variable loss of methylation in old samples and in breast cancer (BC) samples. PAM50 subtypes and estrogen receptor (ER) expression status of the tumors are shown for the BC WGBS tracks. Bottom tracks are RNA-seq tracks from DCIS samples showing cryptic transcription of the HERVH element. All RNA-seq tracks are scaled to the same value and PAM50 subtype is indicated on the right. (B) Heatmap showing signals across 74 TCGA-BRCA samples for ATAC-seq peaks that overlap TEs hypomethylated with age. ATAC-seq peaks (rows) and TCGA BRCA samples (columns) are split into 3 clusters each using k-means clustering. (C) Expression of *ZNF92* in normal tissues and five PAM50 subtypes of breast tumors from the TCGA BRCA collection (p-value Luminal A (LumA) versus Basal and Luminal B (LumB) versus Basal (p-value < 2.2×10^-16^; using two-tailed Wilcoxon signed-rank test).

We next examined whether TEs with methylation loss function as regulatory elements in breast cancer. To this end we utilized ATAC-seq data from The Cancer Genome Atlas (TCGA) breast cancer cohort (TCGA-BRCA) and ChIP-seq datasets from breast cancer cell lines (51). We identified 1197 TCGA-BRCA ATAC-seq peaks that overlap TEs with methylation loss in aging LEps (**Figure 6B**). Although some TEs were accessible in basal subtype tumors (418 of 1197), most of them were accessible across luminal tumors (779 of 1197 peaks) (**Figure 6B**). Some luminal tumors that were diagnosed as lobular carcinomas had particularly high chromatin accessibility across all TEs (**Figure 6B**). These results indicate that TEs hypomethylated with age may function as regulatory elements in breast cancer.

To determine whether loss of methylation at TEs can lead to changes in expression of genes that are thought to be important in breast cancer, we mapped the 554 age-hypomethylated TEs to genes within ±500kb and calculated the correlation of TE expression with each gene mapped within this window in DCIS samples (49). We identified 406 genes that were significantly correlated with the expression of a nearby TE (**Table S8**). A representative example of an L1PA4 element with significant expression correlation (correlation: 0.8, *p*-val: 7.29e^-9^) with the nearby *ZNF92* gene is shown (**Figure S10A**). We find that *ZNF92* is also highly expressed in luminal subtype breast tumors (Luminal A and Luminal B) as compared to basal tumors (**Figure 6C**). These results suggest that TEs located at deleterious locations in the genome are reawakened through stochastic loss of methylation causing expression of genes that are associated with breast cancers. Overall suggesting that reawakening of TEs is directly related to increased cancer susceptibility with age.

## Discussion

Previous studies analyzing aging-associated DNA methylation changes in human mammary luminal epithelial cells have either used array-based technologies (8,11,17,18) or were performed using composite tissues that represented a mixture of cells (17,18). Array probes are mainly enriched at genic regions and intergenic elements including repeats and distal regulatory elements are under-represented (52). It has furthermore been shown that different cell lineages have unique responses to aging (8,53), making pure cell populations critical for interpretation. Our use of WGBS on sorted normal human mammary epithelial cells addresses both shortcomings.

We find that aging leads to DNA methylation changes at lineage-specific gene regulatory elements in mammary luminal epithelial cells, in-line with previous studies that show loss of lineage fidelity in luminal cells with age (8). Specifically, luminal cells show DNA methylation loss at myoepithelial-specific regulatory elements containing MEp-specific TF binding sites. Moreover, luminal-specific regulatory regions containing LEp-specific TF binding sites are repressed by DNA methylation with age. This loss of lineage-fidelity has been shown to occur with age and in mammary tissue that is susceptible to breast cancer (11,54).

Previous DNA methylome studies in human cells suggested a global loss of methylation with age (16,38,55). Those studies were performed using CD4^+^ T cells from a neonate and a centenarian (16), sorted CD4^+^ T cells (38) and skin keratinocytes (55) from young (<25 y) and old (>70 y) individuals. The global methylation loss with age was accompanied by a global increase in entropy or methylation discordance (38). We did not observe a global methylation loss or a global increase in entropy in mammary luminal epithelial cells with age. This could be due to higher fidelity of DNA methylation maintenance in luminal cells or due to a lesser degree of separation between the age of young (<30 years) and old (> 55 years) individuals in our study. DNA methylome studies in aging mouse livers also showed no global loss of methylation, but specific changes at regulatory elements were detected (56). Whereas global methylation loss was not detected, we did detect methylation changes at regulatory elements and variable methylation changes at evolutionarily recent transposons.

Global loss of methylation and focal increase in methylation, termed ‘epigenetic drift’, has been reported to occur with age and in cancer (57). The focal increase in methylation has been reported to occur mainly at CpG islands (CGIs). We find CGIs showing both gain and loss in methylation reflecting changes in lineage-specific gene expression. However, we do observe increased methylation disorder/entropy at CGIs that lead to methylation gains with age, which is further enhanced during the transition to luminal breast cancer. These CGIs were predominantly PRC2 targets that are known to gain methylation in other cancer types (58). PRC2 targets were also found among genes that showed methylation gain at promoters. Loss/repositioning of PRC2 has been proposed to cause DNA methylation gain at its targets. Presumably, CGIs with increased methylation entropy are in the process of becoming hypermethylated in luminal breast cancer. Hypermethylation of PRC2 targets seems to be specific to the luminal subtype of breast cancer (19), although the reasons for this are not currently clear.

DNA methylation is an important mechanism for silencing of transposable elements. Reactivation of transposable elements has been shown in cancer cells, coincident with a global loss of DNA methylation (59,60). Previous research comparing DNA methylation levels of CD4^+^ T cells between a neonate and a centenarian reported that 12% of DMRs that lost methylation with age were located on Alu elements (16), whereas changes in epigenetic silencing at TEs with aging has not been systematically investigated. Increased expression of retrotransposons and active retrotransposition have been reported in aging mouse tissues (61,62) and increased LINE-1 expression has been reported in senescent human fibroblasts *in vitro* (63). However, these studies used bulk quantitative strategies such as RT-qPCR, which are not ideal due to high sequence similarity between multiple active and inactive copies. Our approach allowed us to investigate the impact of aging on DNA methylation at specific TEs using non-senescent mammary epithelial cells. Our results demonstrate that evolutionarily recent (e.g primate and hominid specific) transposons are more likely to be silenced by DNA methylation in young and show increased variability and methylation loss with age. These TEs could be an important component of understanding differential susceptibility to cancer as not all aged individuals develop cancer. Our data also underscore the need for use of human cells or organoids for future studies as these cannot be recapitulated in mice.

We further show that the methylation loss observed at TEs with age becomes exacerbated in frank luminal breast cancers. The derepressed TEs can lead to aberrant transcription or function as regulatory elements in breast cancer cells. We demonstrate that hypomethylated TEs could regulate genes associated with luminal subtype breast cancers. TEs also contribute lineage-specific TF binding sites to regulatory elements associated with loss of lineage fidelity with age. Retrotransposon activation has been widely observed in nearly every human cancer, including breast cancer (64,65), and even in premalignant lesions (60). This suggests that early events in tumorigenesis may be driven by retrotransposon activity. We show here, for the first time, that loss of transposon silencing is evident in aging luminal epithelial cells that could predispose towards breast cancer. Future studies are needed to characterize the role of these TEs in the early events of mammary cancer initiation. Altogether, our results showing DNA methylation alterations at regulatory elements, CGIs and TEs with aging provide further evidence in support of the observation that epigenetic changes with age could determine susceptibility towards breast cancer.

## Materials and Methods

### Isolation of mammary luminal epithelial cells

LEps were isolated from 4^th^ passage (p4) cultures of pre-stasis finite-lifespan HMECs from normal reduction mammoplasty (RM) tissues (6). Primary non-immortalized HMECs were generated and maintained, as described previously (66). HMEC strains were established in the laboratory of Dr. Martha Stampfer (Lawrence Berkeley National Laboratory) who has all required IRB approvals to distribute these samples. Cells were grown in M87A medium with cholera toxin and oxytocin at 0.5 ng/ml and 0.1nM, respectively. 4^th^ passage HMECs were stained with anti-human CD133-PE (Biolegend, clone 7) and anti-human CD271 (Biolegend, clone ME20.4) by following standard flow cytometry protocol. LEps and MEps were enriched using Aria III (Becton Dickinson) flow sorter as CD133^+^/CD271^-^ and CD133^-^/CD271^+^ fractions, respectively. The different HMEC strains used are described in Table S1.

### WGBS

We examined the DNA methylome of primary LEps from younger (< 30y) pre-menopausal and older (> 55y) post-menopausal women. Genomic DNA from FACS enriched LEps were extracted using Quick-DNA Microprep kit (Zymo Research). Whole-genome bisulfite sequencing (WGBS-seq) was performed at Admera Health Biopharma Services. Genomic DNA was bisulfite converted using the Zymo EZ DNA methylation kit (Zymo Research) per manufacturer’s protocol. Library preparation was then performed using Accel-NGS^®^ Methyl-Seq DNA Library kit (Swift Biosciences, Michigan, USA) per manufacturer’s recommendations. Equimolar pooling of libraries was performed based on QC values. Samples were sequenced on an Illumina® NovaSeq S4 (Illumina, California, USA) with a read length configuration of 150bp paired-end. We obtained about 38X coverage per genome to an average of 414 million paired-end reads (Table S2).

Sequencing reads were hard trimmed 15 bp from both ends using Trim Galore (https://github.com/FelixKrueger/TrimGalore) (version 0.4.5) to remove unwanted methylation bias arising from the library preparation. Adapter contamination was also removed using Trim Galore using --illumina option. Trimmed reads were aligned to the human reference genome (hg19) using Bismark (version 0.19.1) (67) with the --bowtie1 option (68). Bismark (deduplicate_bismark) to remove PCR duplicates. Methpipe (69) (version 3.4.3) was used to determine the methylation status of individual CpG sites. The hg19 reference genome contains about 56.4 million CpG loci (hg19) or 28.2 million CpG dyads. Methylation levels were calculated for individual CpG dyads and are henceforth referred to as CpG sites.

### DMR identification

DMRfinder (70) was used to identify DMRs. Briefly, CpG sites with at least 4X coverage in at least 6 out of 10 samples were determined. Genomic regions with at least 4 CpG sites within a distance of 100 bp were identified. The maximum length of regions was set to 500 bp. Of these regions, ones that had less than 20 reads covering them were discarded. The final set of regions was then used to calculate differentially methylated regions with greater than 20% change (*P*-value < 0.05) between pooled young and old samples.

### ATAC-seq and analysis

50,000 live FACS sorted young (*n* = 4) and old (*n* = 4) LEps were used for Tn5 transposition in duplicates using the Omni-ATAC method (71). Libraries were sequenced to a depth of 40 million paired reads each on an Illumina HiSeq 2500 system. Raw sequencing reads were trimmed using Trim Galore (https://github.com/FelixKrueger/TrimGalore) (version 0.4.5) to remove adapters. Trimmed reads were aligned to the human reference genome (hg19) using bowtie (68) with the --best -k 1 -X 2000 -mm --chunkmbs 1024 parameters. Unmapped reads, mates as well as low quality reads and secondary alignments were removed using samtools. PCR duplicates were identified using Picard and removed using samtools. The reads were shifted by 4 bases on the sense strand and 5 bases on the anti-sense strand to account for Tn5 bias. Reads from technical replicates were pooled to generate pseudoreplicates. Peaks were called using MACS2 version 2.1.1.20160309 using -p 0.01 --nomodel --shift −37 --extsize 73 --SPMR --keep-dup all --call-summits as parameters. Peak calls from technical replicates and pseudo-replicates were compared using IDR to get the most conservative set of peaks for each sample.

### Enrichment of DMRs and variably methylated regions at TE subfamilies

The extent of overlap of DMRs and variable regions at repat subfamilies (RepeatMasker annotation) was calculated using bedtools intersect. Randomly shuffled control regions of the same length were generated 1000 times, and the expected number of overlaps calculated using bedtools intersect. Enrichment values were calculated by dividing the number of DMR overlaps (observed) with the average number of random overlaps (expected). *P* values were calculated by dividing the number of extreme values (at either tail of distribution) from random overlaps by 1000. Significantly enriched repeat subfamilies had p < 0.05, enrichment > 1.5-fold, and > 5 copies overlapping DMRs or variable regions.

## Availability of data and materials

All raw and processed sequencing data files have been deposited at NCBI’s Gene Expression Omnibus (GEO) under accession GSE153696 (WGBS and ATAC-seq) and GSE182338 (RNA-seq).

## Authors’ contributions

PS, MAL and DES designed the study. PS, MM and CT performed experiments. PS, RWS, MB and AL analyzed and interpreted the data. PS, MAL and DES prepared the manuscript. All authors read and approved the final manuscript.

## Acknowledgements

We thank other members of the Schones laboratory for helpful discussions and comments, and the continued involvement of patient advocates Susan Samson and Sandy Preto. The results shown and referenced here are based in part upon data generated by the TCGA Research Network: https://www.cancer.gov/tcga, and the ENCODE Consortium: https://www.encodeproject.org/.

