## Supplemental material for "Normal mammary epithelia accumulate DNA methylation changes with age that are consistent with breast cancer methylomes"

### Supplemental Methods

#### Power Analysis

We used previously generated RRBS data from LEps derived from tissue of young (n=6) and old (n=4) women for determining sample size for WGBS. We used MethylSeqDesign [1] to determine sample size needed for different discovery rates. We found that a sample size of 5 per group allows 74.82% power for detecting differences in methylation (FDR < 0.05). A post-hoc analysis of our WGBS data using n=3 as pilot data on chromosome 1 also showed that we have 75% power with n=5 per age group.

#### Characterization of DMRs

Overlaps with genomic features were calculated using bedtools intersect [2] with masked CpG islands from the UCSC genome browser [3] and repeats from the RepeatMasker annotation [4]. DMR overlaps with CpG islands, and repeat elements were determined using bedtools intersect such that the overlapping region contains at least 1 CpG site. Kernel smoothed plots were made using the BSSeq package with default smoothing parameters and CpG sites with greater than 3X coverage [5]. Heatmaps were made using the pheatmap package [6].

#### Identification of variably methylated regions

Variably methylated regions were identified using the Systematic Interindividual Variation (SIV) approach [7]. Briefly, the human genome was divided into 100 bp non-overlapping windows containing at least 1 CpG site. About 14.4 million such windows were obtained. The coverage and methylation values were calculated for each sample across these 100bp windows using Methpipe. Windows were then filtered by coverage as follows to obtain informative bins: Windows with CpG <= 2 that had a coverage of at least 10 reads and windows with CpG > 2 that had a coverage of at least 10 reads at half the number of

CpG sites. We obtained about 12.9 million bins that had sufficient coverage. Next, we calculated the Pearson correlation co-efficient for each bin with its adjacent bin. Adjacent bins that were within 100 bp of each other and a correlation >= 0.71 were merged into correlated blocks of methylation. The average methylation for each sample was calculated for the correlated blocks. Correlated blocks which had an interindividual range (IR) of >= 20% and at least 4 CpG sites were defined as variable regions. Old specific variable regions were defined as regions that had an IR of >= 20% across the five old samples and $IR_{old}$ -$IR_{young}$ >= 20%. Similarly, young specific variable regions were defined as regions that had an IR of >= 20% across the five young samples and $IR_{young}$ -$IR_{old}$ >= 20%. These regions were further filtered to retain those that have coverage of at least 3 CpGs in all 10 samples. To categorize variably methylated regions into variable loss or variable gain, the highest ($old_{max}$) and the lowest ($old_{min}$) methylation values among old samples were compared to the methylation average in young ($avg_y$). The absolute values of differences in methylation between $avg_y$ and $old_{max,}$ $old_{min}$ were compared. If the difference in methylation between the $avg_y$ and $old_{min}$ were higher than that between $avg_y$ and $old_{max}$, that region was assigned as a variable region with methylation loss.

**Chromatin state predictions**

ChIP-seq data for mammary epithelial populations were downloaded using the International Human Epigenome Consortium data portal (https://epigenomesportal.ca/ihec/). Bigwig files containing ChIP-seq signal for six different histone modifications in three cell types (sample IDs: CEMT0154, CEMT0155, and CEMT0156) were used (**Figure S4A-C**). ChIP-seq signals from the bigwig files were binned into 200 bp windows. These binned signal files were then binarized using BinarizeSignal in ChromHMM (version 1.20) [8]. The binarized files were used to run

ChromHMM LearnModel with 16 states. Functionally similar chromatin states were grouped together to seven distinct chromatin states (**Figure S4C**). DMR overlaps with chromatin states were performed using bedtools intersect. In cases where a DMR overlapped more than one chromatin state, DMR was assigned to the state with a higher base overlap.

**RNA-seq analysis**

Total RNA was isolated using Quick-RNA Microprep kit (Zymo Research) from FACS sorted young ($n$ = 10) and old LEps ($n$ = 8). Similarly, total RNA was isolated from young ($n$ = 6) MEps. PolyA-enriched RNA was isolated and used for library preparation using the TruSeq RNA Library Prep kit (Illumina). About 50 million single-end reads of length 51 bp were generated for all samples. Raw sequences were aligned to the hg19 reference genome using HISAT2 (version 2.1.0) [9] with the --dta parameter. Stringtie (version 1.3.4) [10] was used with default parameters to assemble transcripts using the Gencode v19 transcript annotation. Assembled transcripts from all libraries were further merged using --merge option in Stringtie. Merged transcript abundances were measured using bedtools coverage.

For the old LEp versus young LEp comparison, edgeR (v3.26.4) [11] was used to conduct differential gene expression analysis. Briefly, normalization factors were calculated using the trimmed mean of M values method (TMM) to eliminate composition biases between libraries. Genewise biological dispersion was estimated using edgeR's empirical Bayes approach. Then, we employed a generalized linear model (GLM) approach to account for the unnormalized distribution of gene counts, as well as additional factors, such as batch effects. Differentially regulated genes were identified with an absolute log2 fold change $\geq$ 0.5 and FDR < 0.05. Gene expression boxplots were generated using batch corrected

RPKM values. Batch correction of RPKM values was done using limma package [12]. For LEp versus MEp comparison, DESeq2 [13] was used to normalize counts and identify differentially expressed genes (absolute log2 fold change ≥ 0.5 and p-value < 0.05).

RNA-seq data from normal and DCIS samples were obtained from GEO (Accession number: GSE69240 [14]). Fastq reads were quality and adapter trimmed using Trim Galore with the --paired option. Quality trimmed reads were aligned to the hg19 genome using STAR (version 2.6.0.a) [15] with --sjdbOverhang 100 --winAnchorMultimapNmax 100 --outFilterMultimapNmax 100 --outFilterMismatchNmax 3 --alignEndsType EndToEnd options. The SQuIRE pipeline [16] was used to measure gene and TE expression changes in DCIS as compared to normal tissues.

**Motif analysis**

Motif enrichment analysis was performed using findMotifsGenome.pl in homer [20]. For each set of genomic regions, background sequences with matched GC content were selected. *P*-values for motif enrichment was calculated using the cumulative binomial distributions. Motif scanning was performed using fimo [18] with JASPAR 2020 database with default parameters [19]. Only binding sites with TFs that were expressed (TPM > 0.1) in LEps or MEps were used for further analyses.

**Methylation entropy analysis**

Normalized methylation entropy (NME) and mean methylation levels (MML) were calculated on 150bp non-overlapping windows across the genome using informME [22]. Hyper-entropic regions were defined as windows with difference in NME (dNME) > 0.1 between at least 13 out of 25 young and old comparisons and methylation levels > 0.9 or < 0.05 in all young samples.

**Additional data analysis and external data**

Mappability of 100bp bisulfite converted kmers for the hg19 genome generated using Bismap [17] were downloaded from the UCSC Genome Browser [3]. Bedgraph files were used to obtain the mappability score of individual CpGs within TEs. The evolutionary age of TEs was calculated as described previously using the Jukes-Cantor correction [21]. Normalized counts for TCGA-BRCA cancer specific ATAC-seq peaks and RNA-seq counts were downloaded from the Genomic Data Commons (GDC) Portal. The clinical information and PAM50 subtypes for these samples were accessed using TCGAbiolinks [23]. Primary breast tumor WGBS data generated as part of the International Cancer Genome Consortium [24] was obtained from European Genome-phenome Archive after DACO approval. Figures were made in R (version 3.5.1) using the ggplot2 package [25] and assembled in Adobe Illustrator.

### Supplemental Figures

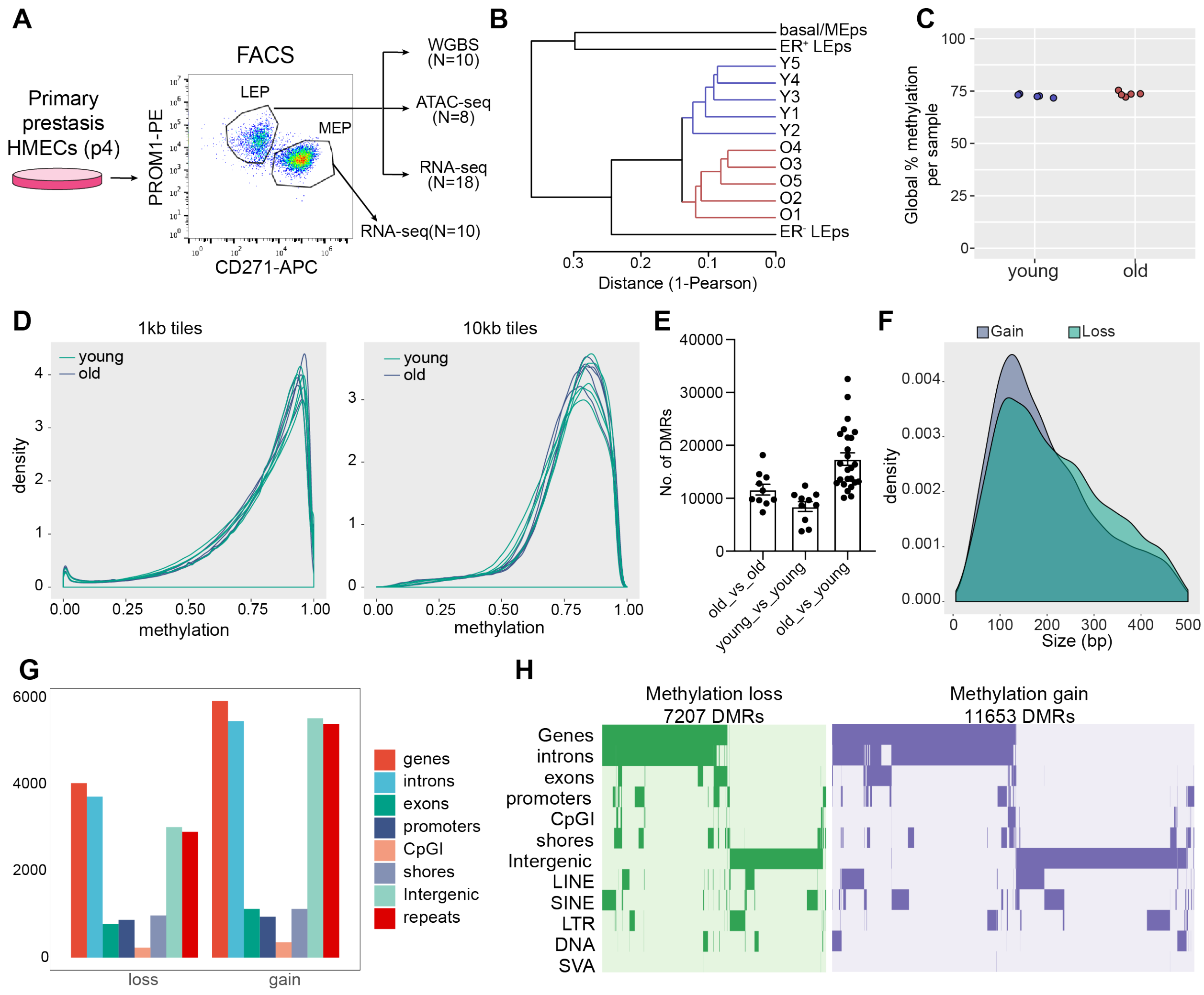


**Figure S1**

**Figure S1: Characterization of whole genome methylation data and DMRs**. (A) Strategy for enrichment of the luminal and myoepithelial cell populations and design of study. (B) Unsupervised hierarchical clustering of young and old LEps and previously published mammary epithelial cell types using methylation levels at all covered CpG sites. (C) Global methylation percentage across all CpG sites in the genome in the ten samples. (D) Density distribution of average DNA methylation values within 1- and 10-kb windows across the genome in young and old LEPs. Only windows with 3 or more CpGs per 1 kb, or 10 or more CpGs per 10 kb, were considered for the analysis. (E) Number of DMRs identified in pairwise comparisons between all samples. (F) Size distribution of DMRs with methylation loss or gain. (G) Number of DMRs with methylation loss or gain overlapping various genomic features as indicated. (H) Overlap of DMRs with a selection of genomic features (rows), including various repeat classes. Features are not exclusive. Green or purple indicates an overlap between a DMR and a feature. DMRs that lose methylation are shown in green, and those that gain methylation are shown in purple.

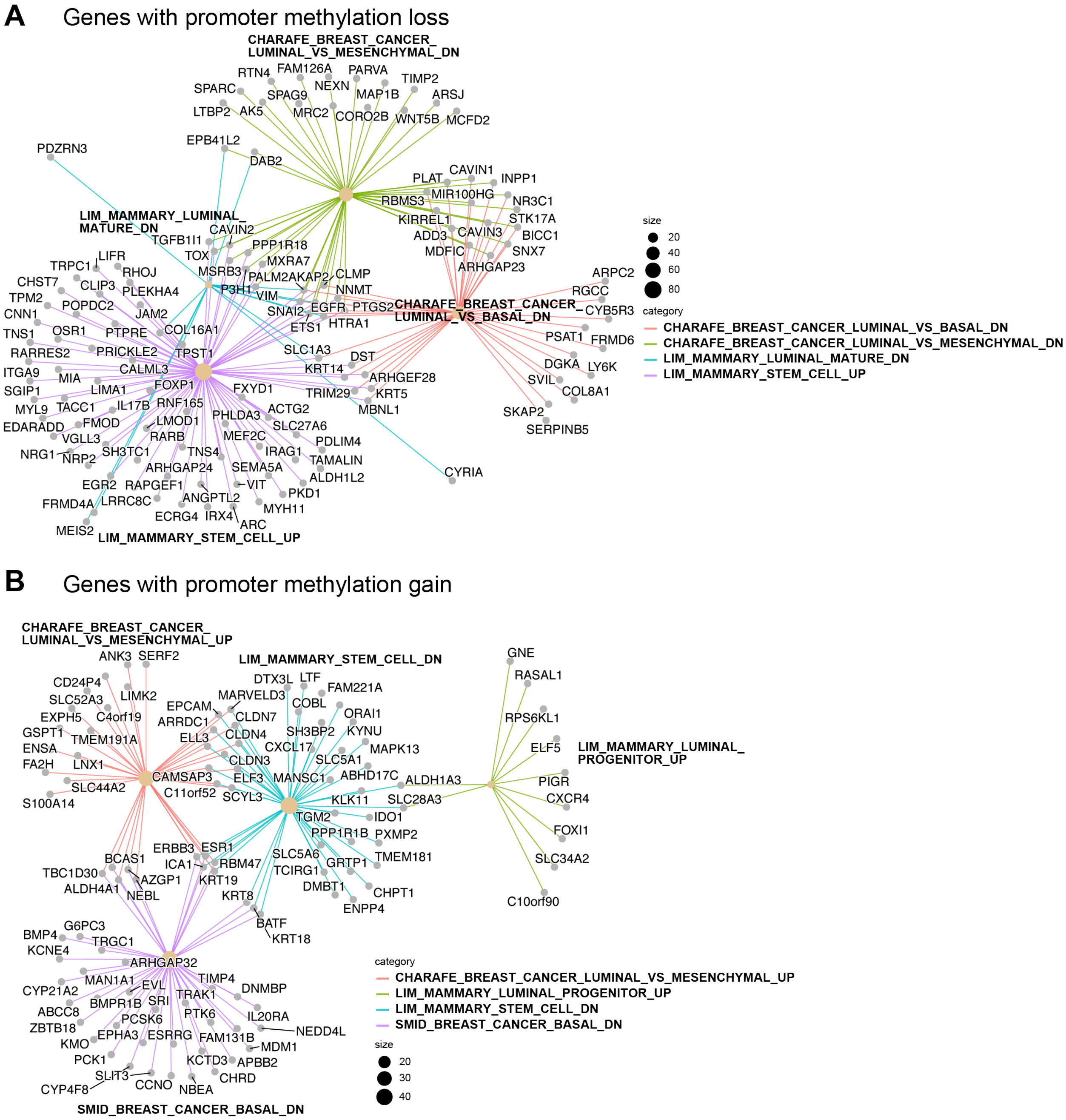


**Figure S2: Lineage-specific genes show methylation changes with age** (A) Genes within select enriched MSigDB gene sets with age dependent DNA methylation loss at promoters. (B) Genes within select enriched MSigDB gene sets with age dependent DNA methylation gain at promoters. Promoters were defined as TSS+/-2kb. UP: upregulated, DN: downregulated.

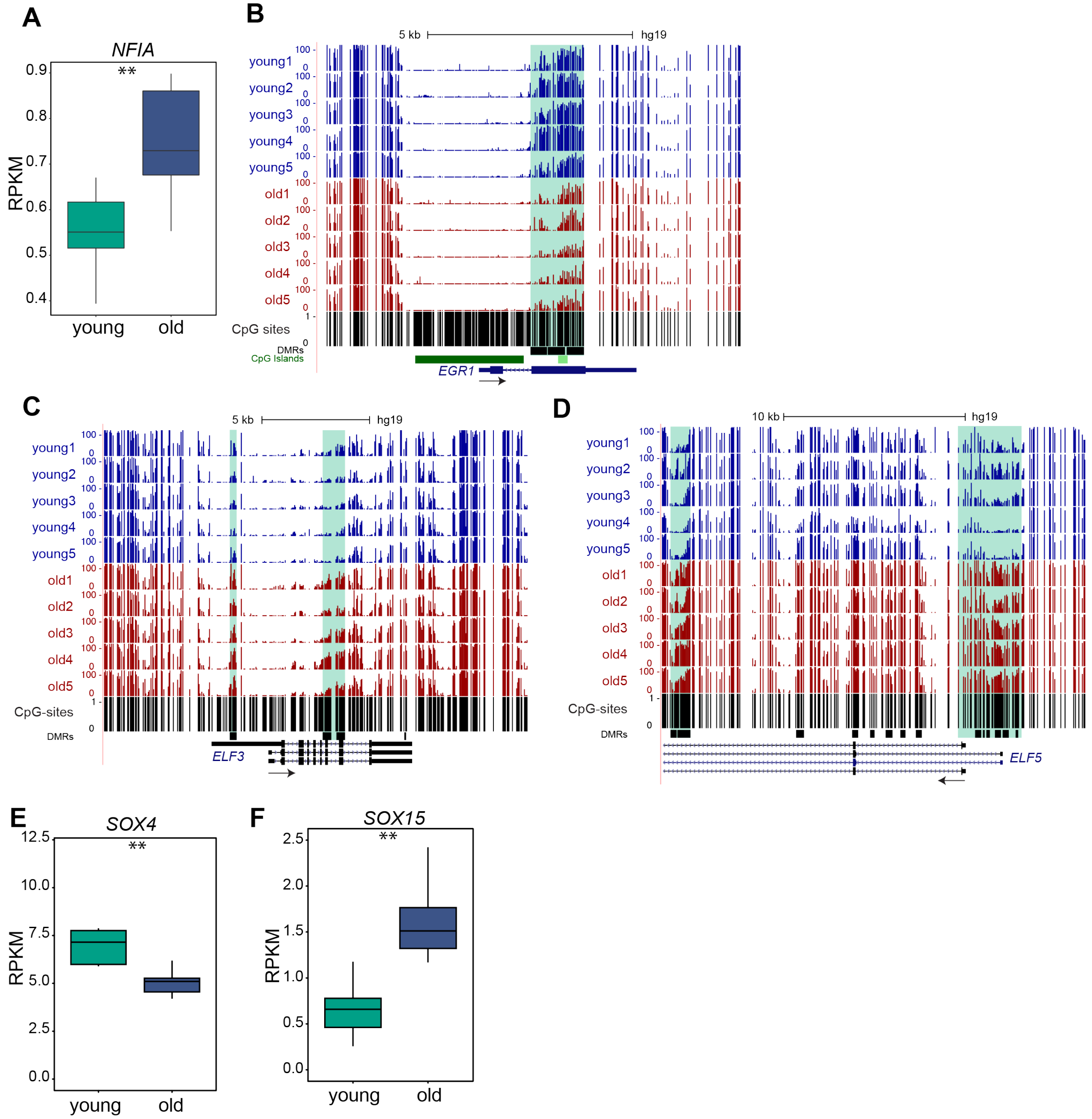


**Figure S3: Lineage-specific transcription factors change expression with age.** (A) Boxplots showing expression levels of *NFIA* in young and old LEPs (B) Genome browser tracks showing DMRs with methylation loss at the *EGR1* gene locus. (C-D) Genome browser tracks showing DMRs with methylation gain at the *ELF3 and ELF5* gene loci. (E-F) Boxplots showing expression levels of transcription factors as indicated in young and old LEPs. For A, E and F; n=8, *p*-value < 0.01; two-tailed unpaired t test with Welch's correction.

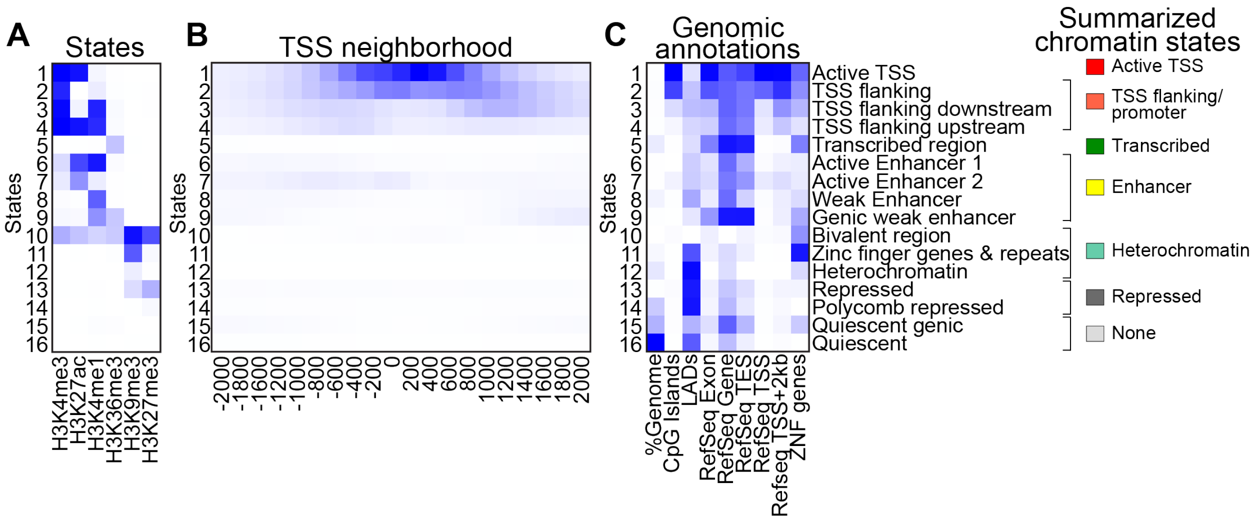


**Figure S4: Chromatin state definitions identified in mammary epithelial lineages.** (A) Chromatin states identified using ChromHMM based on the histone mark probabilities in mammary epithelial lineages. Each row corresponds to a different state, and each column corresponds to a distinct mark for the 16-state model. The darker blue color corresponds to a higher probability of observing the mark in the state. (B) Fold enrichment for each state at 200-bp bin position within 2 kb around a set of transcription start sites (TSSs). A darker blue color corresponds to a higher fold enrichment. (C) Fold enrichment for overlap of each state with various genomic features as indicated. A darker blue color corresponds to a higher fold enrichment. Right most panel indicates the summarized 7 distinct chromatin states which were generated by combining similar states as indicated in the fugure.

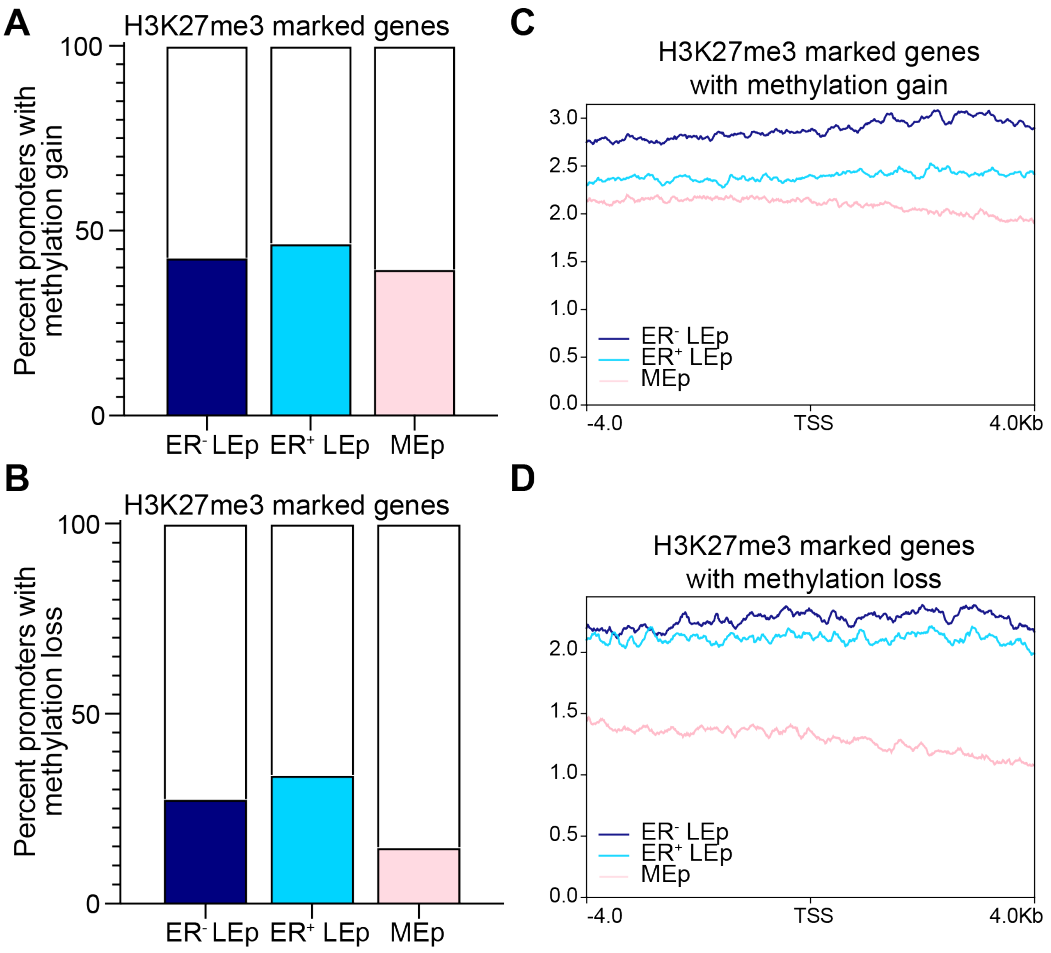


**Figure S5: PRC2 target genes gain promoter methylation in old LEps.** (A) Percentage number of promoters with DNA methylation gain that overlap H3K27me3 peaks (filled bar) in $ER^-$, $ER^+$ luminal cells, and basal cells is shown. (B) Percentage number of promoters with DNA methylation gain that overlap H3K27me3 peaks (filled bar) in indicated mammary cell types. (C) Aggregate plot of H3K27me3 signal around the TSS of genes with age-dependent DNA methylation gain and marked with H3K27me3 in all three cell types (union set). (D) Aggregate plot of H3K27me3 signal around the TSS of genes with age-dependent DNA methylation loss and marked with H3K27me3 in all three cell types (union set).

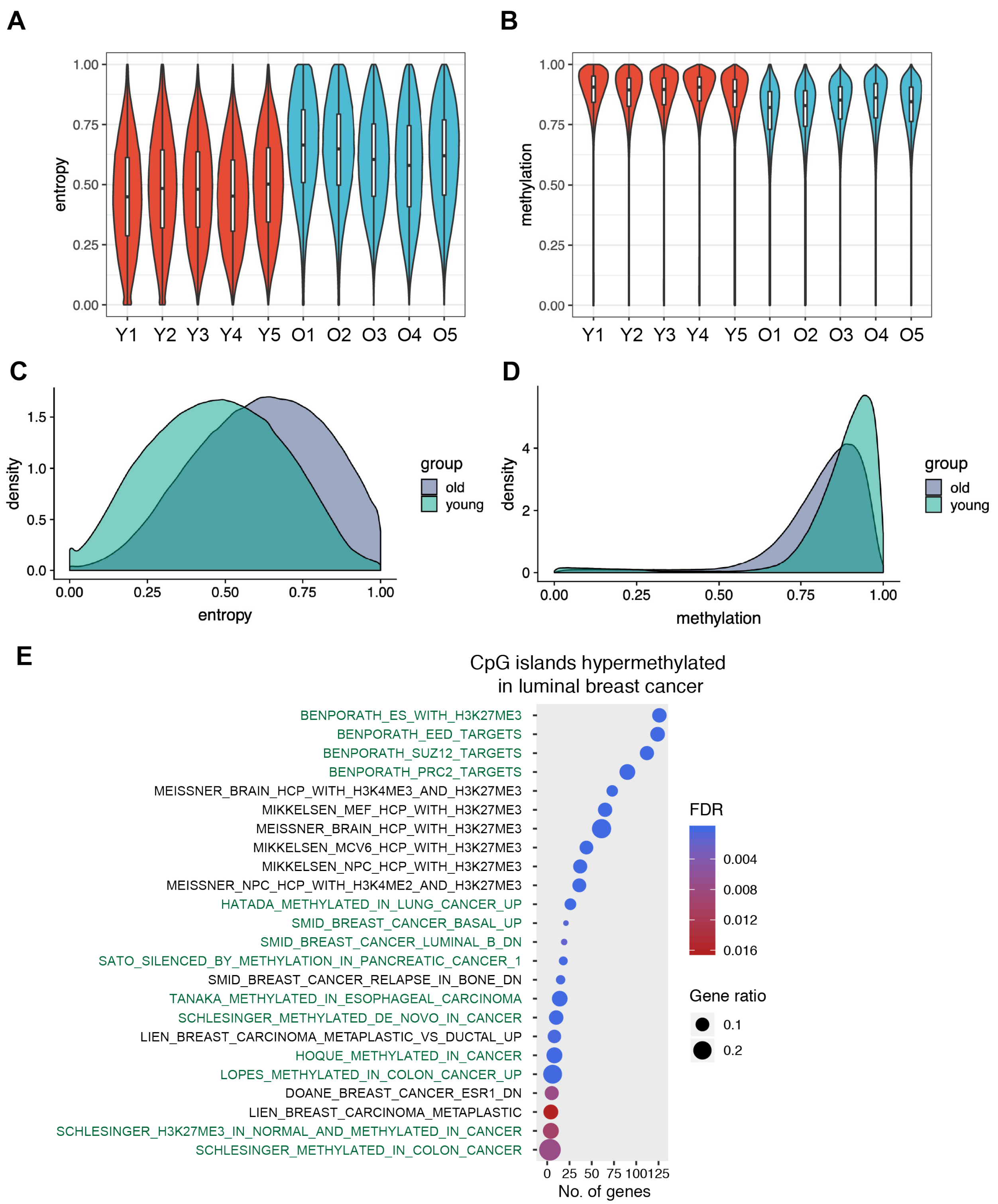


**Figure S6: Regions showing increased methylation entropy in older individuals.** (A) Normalized methylation entropy (Entropy) levels and (B) methylation levels at regions with increased methylation entropy in old individuals. (C) Density plots showing entropy and (D) methylation levels at regions with increased methylation entropy. (E) MSigDB gene sets enriched in CpG island promoters that gained methylation entropy with age and were hypermethylated in luminal breast cancer. Gene ratio denotes the proportion of genes represented out of the total genes in gene set. Relevant gene sets are highlighted in green. BENPORATH gene sets represent PRC2 target genes identified in embryonic stem cells.

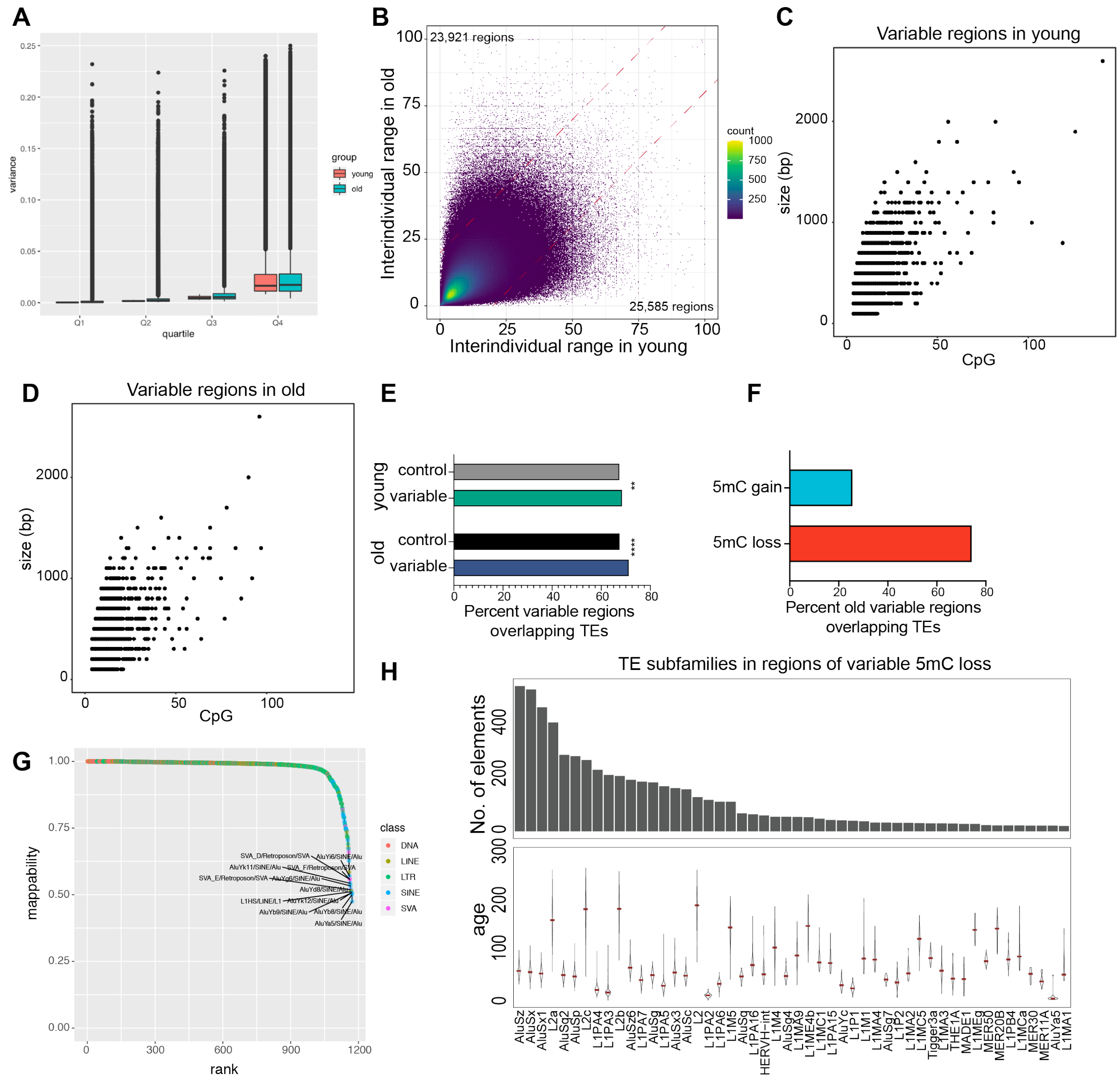


**Figure S7: Characterization of variably methylated regions in luminal cells.** (A) Boxplot shows variance of CpG methylation among young and old samples stratified by variance quartiles in young samples. (B) Plot shows interindividual range of correlated methylation blocks in young and old samples. Points are binned to show density as indicated by color scale. (C) Scatterplot shows the genomic size of the young-specific variable regions versus no. of CpGs in the region. (D) Same as B but showing variable regions specific to young. (E) Percent of variable regions in young and old that overlap TEs. Overlap of shuffled control regions with TEs is shown. *P*-value was calculated by comparing with control regions using Fisher's exact test. **** *p-val* < 0.0001 and ** *p-val* < 0.01. (F) Percent of variable TE regions in old that lose or gain methylation. (G) Average mappability of 100bp bisufite converted reads within TE subfamilies. Indicated subfamilies have average mappability < 0.6. Mappability of 1 denotes unique mapping; 0.5

denotes multi-mapping at 2 sites. (H) TE subfamilies enriched in regions with variable methylation loss. Panels show evolutionary age (top) and number of elements from each subfamily that lose methylation (bottom). In the evolutionary age panels, brown points indicate the median.

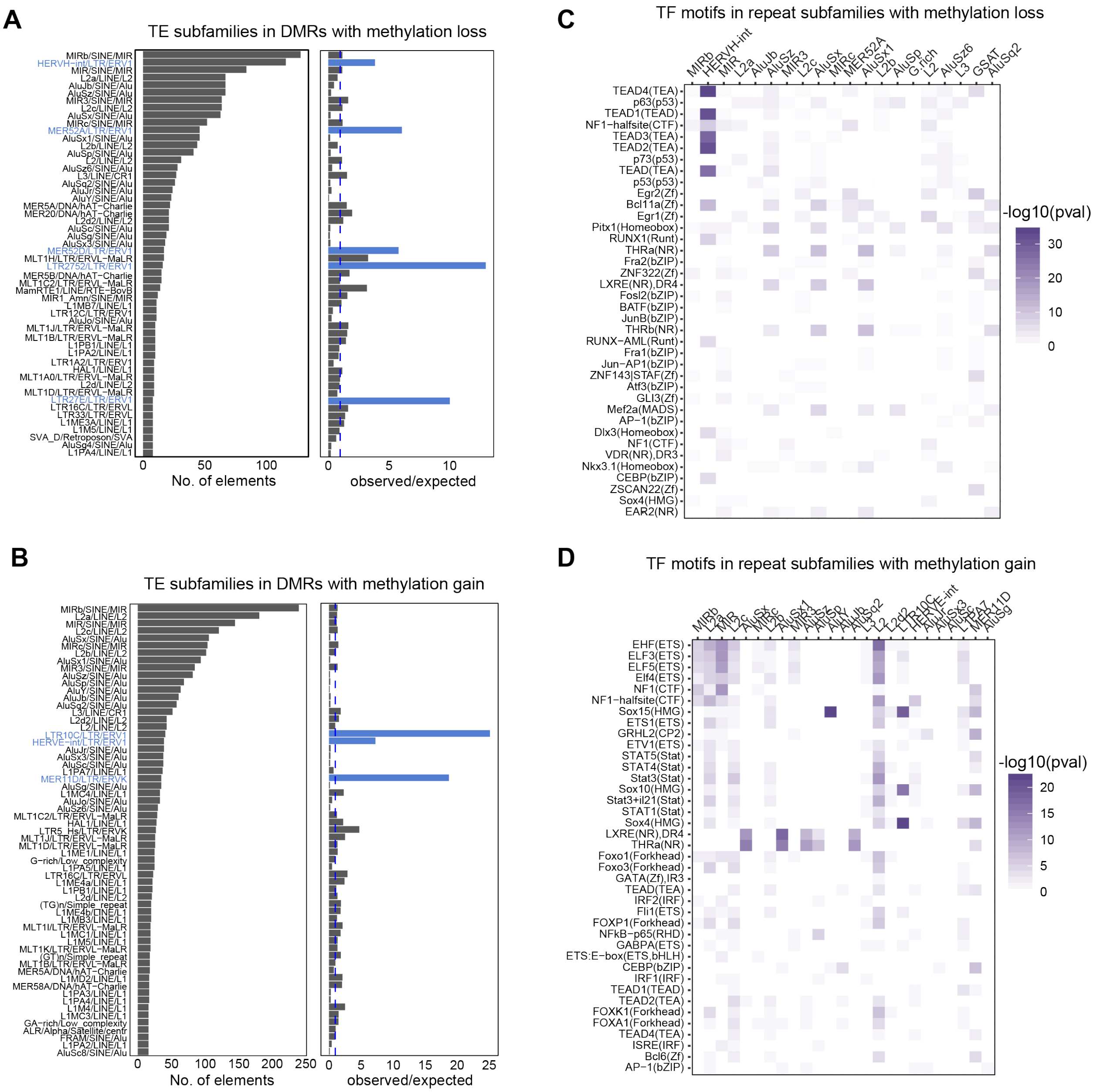


**Figure S8: Transposable elements contribute lineage-specific TF binding sites at DMR regions associated with loss of lineage fidelity.** (A) TE subfamilies enriched in DMRs with methylation loss and (B) methylation gain. A-B Left panel shows number of elements from each subfamily that lose methylation. Right panel shows the observed/expected ratio for each TE subfamily. Expected number of overlaps in a background set of regions with matched length and CpG density was used to calculate the ratio. (C) Enrichment of TF binding site in each TE subfamily with methylation loss and (D) methylation gain. C-D Color scale is –log10(p-value) of TF motif enrichment.

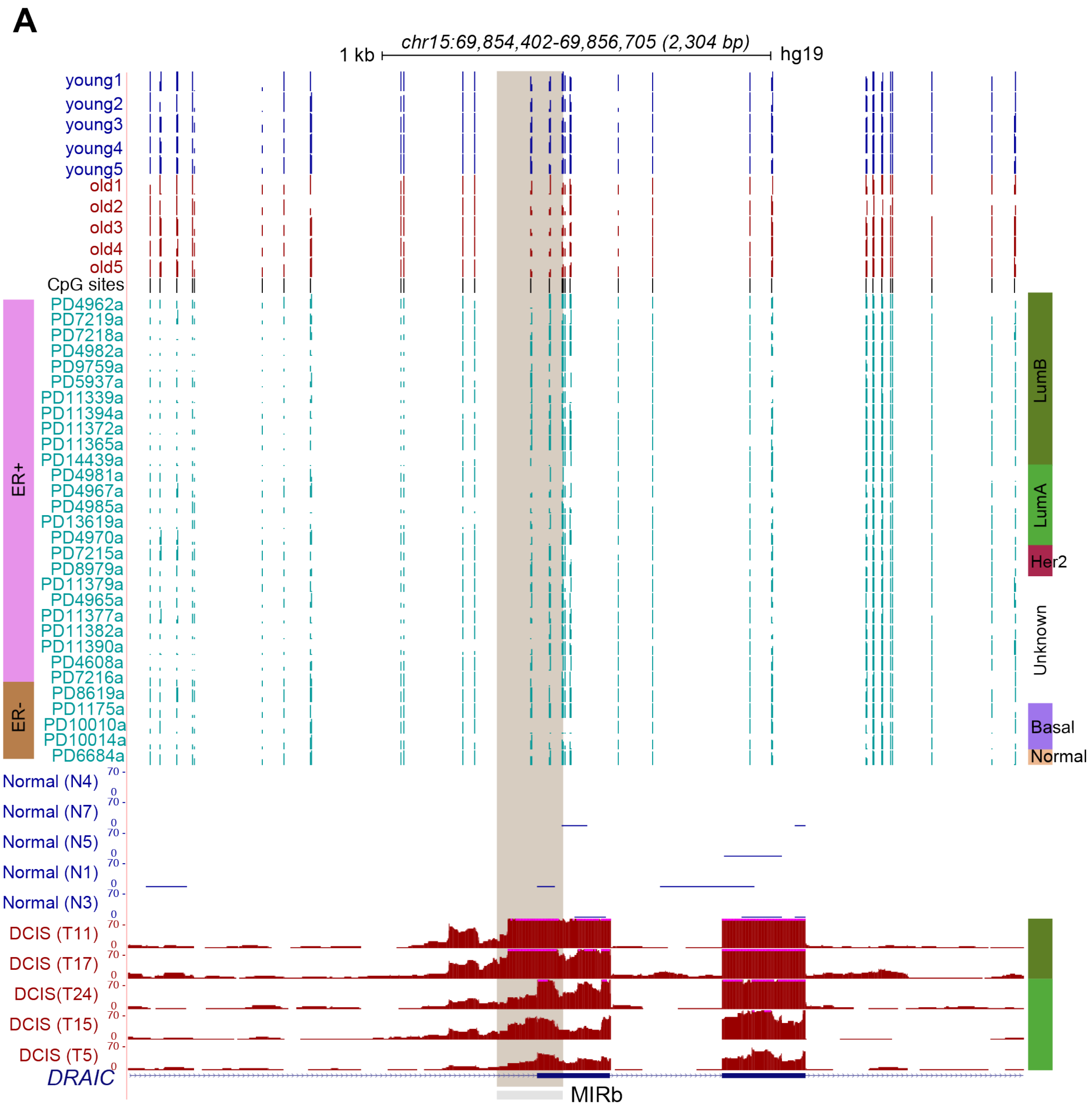


**Figure S9: TEs are activated in DCIS samples.** (A) A representative MIRb element showing methylation loss in old samples and in breast cancer (BC) samples. Bottom tracks are RNA-seq tracks from DCIS samples showing exonization of the MIRb element. All RNA-seq tracks are scaled to the same value.

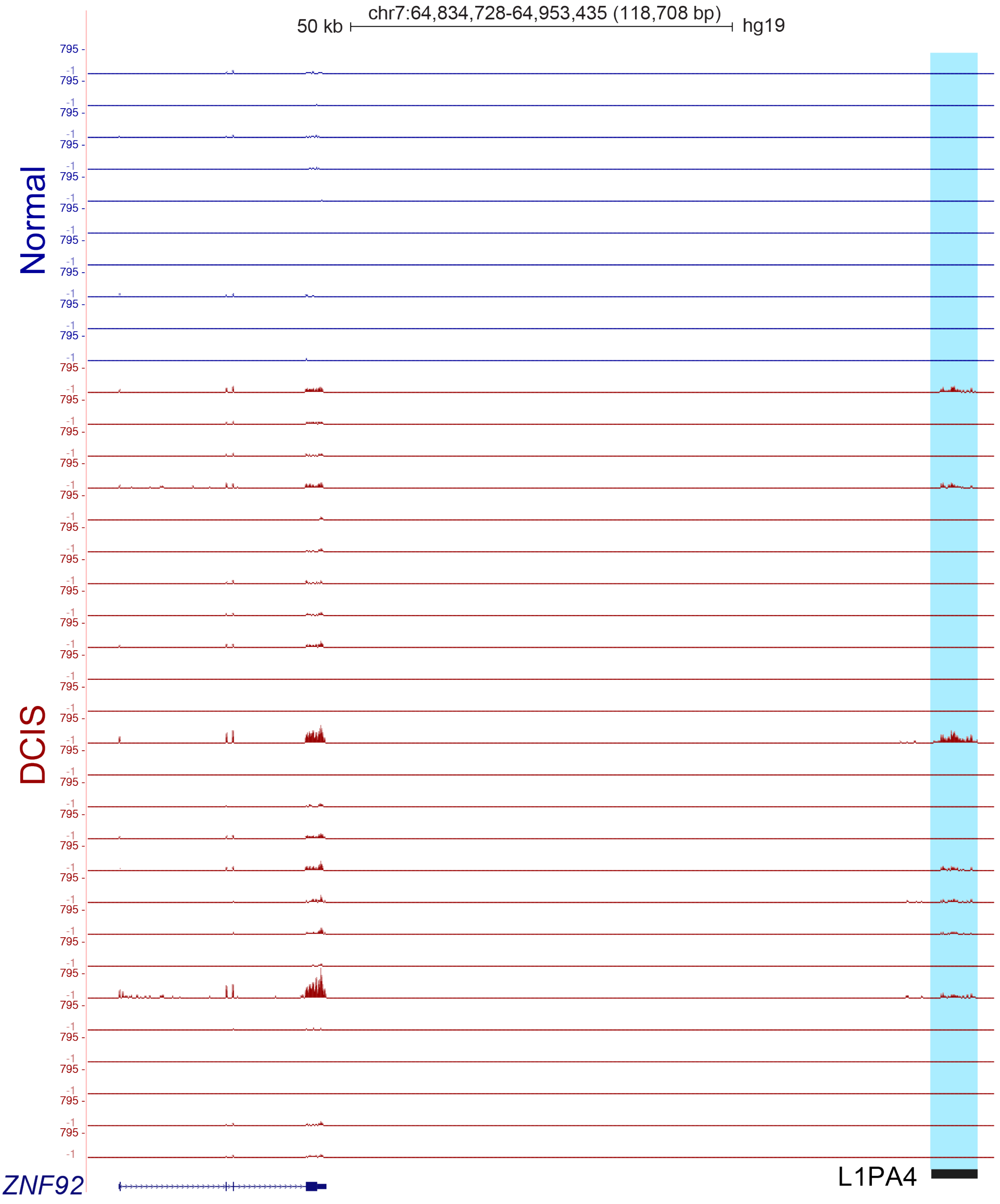


**Figure S10: TEs regulate expression of nearby genes expressed in DCIS and luminal breast cancer.** (A) RNA-seq tracks showing correlated expression of an L1PA4 element and a nearby ZNF92 gene. All tracks are scaled to the same value.

### Supplemental Tables

| Sample name | Age | Sequencing | Sample IDs |
|---|---|---|---|
| Young1 | 19 | WGBS & ATAC-seq | 240L |
| Young2 | 28 | WGBS & ATAC-seq | 172L |
| Young3 | 27 | WGBS | 51L |
| Young4 | 29 | WGBS | 124 |
| Young5 | 21 | WGBS | 356E |
| Young6 | 16 | ATAC-seq | 160 |
| Young7 | 19 | ATAC-seq | 168R |
| Old1 | 56 | WGBS & ATAC-seq | 117R |
| Old2 | 61 | WGBS & ATAC-seq | 112R |
| Old3 | 66 | WGBS & ATAC-seq | 237 |
| Old4 | 68 | WGBS | 029 |
| Old5 | 72 | WGBS | 429ER |
| Old6 | 56 | ATAC-seq | 191L |

**Table S1: Information about HMEC strains used in this study.** RM: Reduction Mammoplasty

| Dataset | Total PE Reads | Uniquely aligned | Unaligned | Non-unique | Mapping Efficiency | % meth in CpG | % meth in CHG | % meth in CHH |
|---|---|---|---|---|---|---|---|---|
| young1 | 442,374,010 | 333,575,143 | 84,149,098 | 24,649,769 | 75.4% | 72.20% | 0.80% | 0.90% |
| young2 | 377,285,149 | 284,644,571 | 69,643,612 | 22,996,966 | 75.4% | 71.70% | 0.90% | 1.00% |
| old1 | 447,412,429 | 331,566,355 | 99,511,987 | 16,334,087 | 74.1% | 72.30% | 1.10% | 1.20% |
| old2 | 389,294,496 | 294,743,268 | 70,389,174 | 24,162,054 | 75.7% | 71.20% | 0.80% | 1.00% |
| young3 | 540,610,191 | 408,246,293 | 116,022,462 | 16,341,436 | 74.8% | 72.9% | 0.5% | 0.5% |
| young4 | 438,076,546 | 267,678,351 | 162,876,069 | 7,522,126 | 73.1% | 72.8% | 0.6% | 0.6% |
| young5 | 678,066,097 | 416,805,644 | 246,854,983 | 14,405,470 | 74.0% | 71.4% | 0.6% | 0.7% |
| old3 | 621,427,696 | 464,255,562 | 142,515,781 | 14,656,353 | 74.7% | 72.9% | 0.5% | 0.4% |
| old4 | 570,538,865 | 416,681,703 | 141,544,457 | 12,312,705 | 73.0% | 74.6% | 0.5% | 0.5% |
| old5 | 477,291,517 | 347,182,922 | 119,705,108 | 10,403,487 | 72.7% | 72.8% | 0.5% | 0.5% |

### Table S2: WGBS alignment statistics

| Dataset | Total CpG sites covered | Fraction covered | Mean depth | Mean depth at covered sites | Mean methylation | Weighted mean methylation | Fractional methylation |
|---|---|---|---|---|---|---|---|
| young1 | 25846312 | 0.92 | 9.69 | 10.58 | 0.74 | 0.73 | 0.8 |
| young2 | 25677175 | 0.91 | 8.95 | 9.84 | 0.74 | 0.73 | 0.8 |
| old1 | 25561113 | 0.91 | 7.33 | 8.09 | 0.74 | 0.73 | 0.79 |
| old2 | 25455720 | 0.9 | 8.2 | 9.09 | 0.74 | 0.72 | 0.8 |
| young3 | 26967650 | 0.96 | 9.48 | 9.94 | 0.747 | 0.74 | 0.80 |
| young4 | 27220228 | 0.97 | 18.56 | 19.24 | 0.744622 | 0.74 | 0.80 |
| young5 | 27267663 | 0.97 | 24.45 | 25.30 | 0.734071 | 0.72 | 0.79 |
| old3 | 27283872 | 0.97 | 27.86 | 28.81 | 0.749737 | 0.74 | 0.81 |
| old4 | 27272794 | 0.97 | 23.78 | 24.60 | 0.76192 | 0.75 | 0.82 |
| old5 | 27255122 | 0.97 | 19.96 | 20.67 | 0.7438 | 0.74 | 0.81 |

### Table S3: WGBS CpG methylation statistics

### Table S4-S6: Attached as excel files

| Top 20 motifs enriched at DMRs with methylation loss | | | | | |
|---|---|---|---|---|---|
| **Motif Name** | **Consensus** | **P-value** | **Log P-value** | **q-value (Benjamini)** | **Sensitivity to DNA methylation (Yin et al, Science, 2017)** |
| Egr1(Zf) | TGCGTGGGYG | 1.00E-91 | -2.11E+02 | 0 | Unaffected |

| | | | | | |
|---|---|---|---|---|---|
| Egr2(Zf) | NGCGTGGGCGGR | 1.00E-75 | -1.74E+02 | 0 | Unaffected |
| p63(p53) | NNDRCATGYCYNRRCATGYH | 1.00E-72 | -1.67E+02 | 0 | No CpG in binding site |
| NF1-halfsite(CTF) | YTGCCAAG | 1.00E-72 | -1.67E+02 | 0 | No CpG in binding site |
| p73(p53) | NRRRCAWGTCCDGRCATGYY | 1.00E-55 | -1.29E+02 | 0 | No CpG in binding site |
| p53(p53) | RRCATGYCYRGRCATGYYYN | 1.00E-53 | -1.23E+02 | 0 | No CpG in binding site |
| p53(p53) | RRCATGYCYRGRCATGYYYN | 1.00E-53 | -1.23E+02 | 0 | No CpG in binding site |
| TEAD4(TEA) | CCWGGAATGY | 1.00E-28 | -6.66E+01 | 0 | No CpG in binding site |
| RUNX1(Runt) | AAACCACARM | 1.00E-28 | -6.59E+01 | 0 | No CpG in binding site |
| TEAD3(TEA) | TRCATTCCAG | 1.00E-27 | -6.38E+01 | 0 | No CpG in binding site |
| TEAD1(TEAD) | CYRCATTCCA | 1.00E-24 | -5.64E+01 | 0 | No CpG in binding site |
| RUNX(Runt) | SAAACCACAG | 1.00E-22 | -5.13E+01 | 0 | No CpG in binding site |
| Lhx2(Homeobox) | TAATTAGN | 1.00E-20 | -4.64E+01 | 0 | No CpG in binding site |
| TEAD(TEA) | YCWGGAATGY | 1.00E-19 | -4.51E+01 | 0 | No CpG in binding site |
| Tlx?(NR) | CTGGCAGSCTGCCA | 1.00E-18 | -4.36E+01 | 0 | No CpG in binding site |
| Dlx3(Homeobox) | NDGTAATTAC | 1.00E-17 | -4.04E+01 | 0 | No CpG in binding site |
| Lhx3(Homeobox) | ADBTAATTAR | 1.00E-17 | -3.99E+01 | 0 | No CpG in binding site |
| Fra1(bZIP) | NNATGASTCATH | 1.00E-16 | -3.80E+01 | 0 | No CpG in binding site |
| Lhx1(Homeobox) | NNYTAATTAR | 1.00E-15 | -3.62E+01 | 0 | No CpG in binding site |
| Fra2(bZIP) | GGATGACTCATC | 1.00E-15 | -3.54E+01 | 0 | No CpG in binding site |
| **Top 20 motifs enriched at DMRs with methylation gain** | | | | | |
| **Motif Name** | **Consensus** | **P-value** | **Log P-value** | **q-value (Benjamini)** | **Sensitivity to DNA methylation (Yin et al, Science, 2017)** |
| EHF(ETS) | AVCAGGAAGT | 1.00E-256 | -5.90E+02 | 0 | Unknown |
| ELF3(ETS) | ANCAGGAAGT | 1.00E-214 | -4.94E+02 | 0 | MethylMinus: sensitive to DNA methylation |
| ELF5(ETS) | ACVAGGAAGT | 1.00E-185 | -4.27E+02 | 0 | MethylMinus: sensitive to DNA methylation |
| Elf4(ETS) | ACTTCCKGKT | 1.00E-173 | -4.00E+02 | 0 | MethylMinus: sensitive to DNA methylation |
| ERG(ETS) | ACAGGAAGTG | 1.00E-134 | -3.09E+02 | 0 | MethylMinus: sensitive to DNA methylation |
| ETV1(ETS) | AACCGGAAGT | 1.00E-101 | -2.33E+02 | 0 | MethylMinus: sensitive to DNA methylation |
| NF1-halfsite(CTF) | YTGCCAAG | 1.00E-94 | -2.19E+02 | 0 | No CpG in binding site |
| Etv2(ETS) | NNAYTTCCTGHN | 1.00E-85 | -1.96E+02 | 0 | MethylMinus: sensitive to DNA methylation |
| ETS1(ETS) | ACAGGAAGTG | 1.00E-72 | -1.66E+02 | 0 | Unknown |
| ETV4(ETS) | ACCGGAAGTG | 1.00E-68 | -1.59E+02 | 0 | MethylMinus: sensitive to DNA methylation |
| GABPA(ETS) | RACCGGAAGT | 1.00E-67 | -1.56E+02 | 0 | MethylMinus: sensitive to DNA methylation |

| | | | | | |
|---|---|---|---|---|---|
| EWS:ERG-fusion(ETS) | ATTTCCTGTN | 1.00E-62 | -1.45E+02 | 0 | MethylMinus: sensitive to DNA methylation |
| NF1(CTF) | CYTGGCABNSTGCCAR | 1.00E-60 | -1.39E+02 | 0 | No CpG in binding site |
| Fli1(ETS) | NRYTTCCGGH | 1.00E-57 | -1.34E+02 | 0 | MethylMinus: sensitive to DNA methylation |
| SPDEF(ETS) | ASWTCCTGBT | 1.00E-55 | -1.27E+02 | 0 | MethylMinus: sensitive to DNA methylation |
| ELF1(ETS) | AVCCGGAAGT | 1.00E-46 | -1.08E+02 | 0 | MethylMinus: sensitive to DNA methylation |
| Stat3+il21(Stat) | SVYTTCCNGGAARB | 1.00E-45 | -1.04E+02 | 0 | Unknown |
| Foxo1(Forkhead) | CTGTTTAC | 1.00E-43 | -1.01E+02 | 0 | No CpG in binding site |
| AP-2alpha(AP2) | ATGCCCTGAGGC | 1.00E-43 | -1.01E+02 | 0 | MethylMinus: sensitive to DNA methylation |
| Tlx?(NR) | CTGGCAGSCTGCCA | 1.00E-43 | -9.97E+01 | 0 | No CpG in binding site |

**Table S7: Known binding sensitivities for TFs with sites enriched at DMRs with methylation loss and gain**
